# Chromatin regulators in the TBX1 network confer risk for conotruncal heart defects in 22q11.2DS and sporadic congenital heart disease

**DOI:** 10.1101/2022.09.30.507111

**Authors:** Yingjie Zhao, Yujue Wang, Lijie Shi, Donna M. McDonald-McGinn, T. Blaine Crowley, Daniel E. McGinn, Oanh T. Tran, Daniella Miller, Elaine Zackai, H. Richard Johnston, Eva W. C. Chow, Jacob A.S. Vorstman, Claudia Vingerhoets, Therese van Amelsvoort, Doron Gothelf, Ann Swillen, Jeroen Breckpot, Joris R. Vermeesch, Stephan Eliez, Maude Schneider, Marianne B.M. van den Bree, Michael J. Owen, Wendy R. Kates, Gabriela M. Repetto, Vandana Shashi, Kelly Schoch, Carrie E. Bearden, M. Cristina Digilio, Marta Unolt, Carolina Putotto, Bruno Marino, Maria Pontillo, Marco Armando, Stefano Vicari, Kathleen Angkustsiri, Linda Campbell, Tiffany Busa, Damian Heine-Suñer, Kieran C. Murphy, Declan Murphy, Sixto García-Miñaúr, Luis Fernández, International 22q11.2 Brain and Behavior Consortium, Elizabeth Goldmuntz, Raquel E. Gur, Beverly S. Emanuel, Deyou Zheng, Christian R. Marshall, Anne S. Bassett, Tao Wang, Bernice E. Morrow

## Abstract

**Background:** Congenital heart disease (CHD) affecting the conotruncal region of the heart, occur in half of patients with 22q11.2 deletion syndrome. This syndrome is a rare disorder with relative genetic homogeneity that can facilitate identification of genetic modifiers. Haploinsufficiency of *TBX1*, mapped to the 22q11.2 region, encoding a T-box transcription factor, is one of the main genes for the etiology of the syndrome. We suggest that genetic modifiers of CHD in patients with 22q11.2 deletion syndrome may be in the *TBX1* gene network.

**Methods:** To identify genetic modifiers of 22q11.2 deletion syndrome, we analyzed whole genome sequence of subjects with 22q11.2DS, of which 456 were cases with conotruncal heart defects and 537 were controls with normal cardiac structures. We retained the most damaging rare coding variants and examined 19 functional gene sets for association that were weighted upon expression of genes in cardiac progenitor cells in mouse embryos identified by RNA-sequencing.

**Results:** We identified rare damaging coding variants in chromatin regulatory genes as modifiers of conotruncal heart defects in 22q11.2DS. Chromatin genes with recurrent damaging variants include *EP400, KAT6A, KMT2C, KMT2D, NSD1, CHD7* and *PHF21A*. In total, we identified 37 chromatin regulatory genes, that may increase risk for conotruncal heart defects in 8.5% of 22q11.2 deletion syndrome cases. Many of these genes were identified as risk factors for sporadic CHD in the general population increasing the likelihood that these genes are medically important contributors for CHD. These genes are co-expressed in cardiac progenitor cells with *TBX1*, suggesting that they may be in the same genetic network. Some of the genes identified, such as *KAT6A, KMT2C, CHD7* and *EZH2*, have been previously shown to genetically interact with *TBX1* in mouse models, providing mechanistic validation of these genes found.

**Conclusions:** Our findings indicate that disturbance of chromatin regulatory genes impact a *TBX1* gene network serving as genetic modifiers of 22q11.2 deletion syndrome. Since some of these chromatin regulatory genes were found in individuals with sporadic CHD, we suggest that there are shared mechanisms involving the *TBX1* gene network in the etiology of CHD.

## Introduction

Congenital heart disease (CHD) occurs sporadically in approximately 1% of the general population resulting in significant morbidity and mortality. A subset has syndromic causes with known genetic etiologies. The 22q11.2 deletion syndrome (22q11.2DS; also named DiGeorge syndrome or velo-cardio-facial syndrome) is an example of a genetic syndrome in which the majority have CHD. It is estimated that ~ 40% of individuals with 22q11.2DS have conotruncal heart defects (CTDs) affecting the formation of the cardiac outflow tract (OFT) and/or aortic arch (1). *TBX1* encodes a T-box transcription factor mapping to the hemizygously deleted region on 22q11.2 and it has been shown to be largely responsible for the syndrome’s cardiac phenotype (2–4). Further, mutation of one allele of *TBX1*, without a 22q11.2 deletion, is responsible for CTDs, confirming its importance as a disease gene (5). Inactivation of one allele of *Tbx1* in mice results in mild aortic arch anomalies while inactivation of both alleles results in a persistent truncus arteriosus, with complete penetrance (2–4, 6). Although haploinsufficiency of *TBX1* has a major impact in the etiology of disease, it cannot fully explain variation in the frequency of occurrence of CTDs and therefore it is likely that this variation is due to the existence of genetic modifiers.

Our goal is to identify genes that modify the effects of the 22q11.2 deletion as related to the haploinsufficiency of *TBX1*. One possibility is that there are damaging coding/splicing variants on the remaining allele of *TBX1* or other genes in the hemizygous 22q11.2 region that can alter risk of CTDs, however mutations were not detected (7, 8). Therefore, we decided to investigate potentially damaging rare coding/splicing variants in the rest of the genome that may serve as modifiers of 22q11.2DS that might be in the *TBX1* genetic network.

In this report, we analyzed whole genome sequence (WGS) of 1,182 subjects with 22q11.2DS, where we focused upon 456 cases with CTDs versus 537 with a normal heart serving as controls to identify variants that affect protein function. We identified specific chromatin regulatory genes including histone lysine acetyltransferases and lysine methyltransferases, of which some are in the *TBX1* molecular network, and secondly, that they strongly overlap with genes found as risk factors for sporadic CHD in the general population.

## Methods

### Study cohort and definition of case vs control groups

A total of 1,595 subjects with 22q11.2DS were recruited by the International 22q11.2 Brain and Behavior Consortium (IBBC); details on ascertainment and original study design were provided previously (8–11). We adopted the same definitions of CHD and CTD cases within, and controls as described in detail in our previous study (8). Briefly, participants with an intracardiac and/or aortic arch defect were considered as a CHD case. Individual diagnoses were obtained within the CHD case category. Given high prevalence in the general population (12, 13), individuals with minor heart anomalies (VSD, ASD, or persisting foramen ovale that closed spontaneously in infancy), or with bicuspid aortic valve, in the absence of other malformations, were considered to be controls.

Of the 1,595 22q11.2DS samples sequenced, 1,182 passed the quality control (QC) procedures for the current study (Fig. 1A, Additional file 2: Figure S1). Briefly, samples with no CHD phenotype information, and those with a 22q11.2 deletion other than the typical 3 million base pair LCR22A-LCR22D deletion were removed. Contaminated and mixed samples as determined by Identity-by-status based on common and independent variants were removed. Only one of related sample pairs and duplicated sample pairs was retained for analysis. After sample QC measures, the cohort consisted of 537 controls, and 645 CHD cases among which 456 have a CTD.

**Fig. 1.**
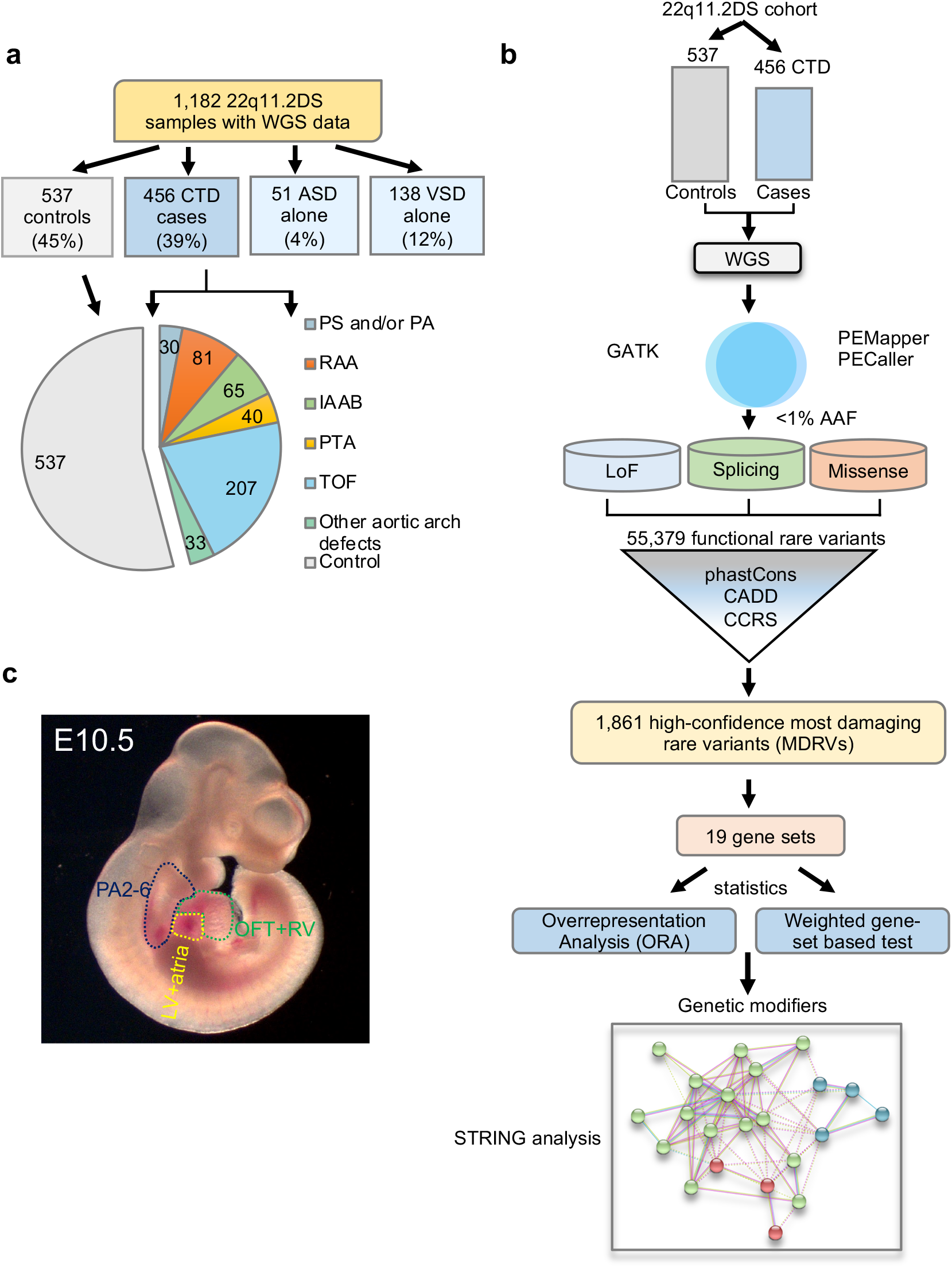
The 22q11.2 deletion syndrome cohort and study design. A, Pie chart of intracardiac and aortic arch phenotypes. Control (gray, no significant heart defect); CTD (conotruncal heart defect, blue); ASD alone (isolated atrial septal defect, light blue); VSD alone (isolated ventricular septal defect, light blue). Pie chart includes controls (gray) versus CTD cases with phenotypes including: TOF (tetralogy of Fallot, light blue), RAA (right sided aortic arch, orange), IAAB (interrupted aortic arch type B, green), PTA (persistent truncus arteriosus, yellow), PS/PA (pulmonary stenosis and/or pulmonic atresia, blue) and other aortic arch defects such as abnormal origin of the right or left subclavian artery, alone (light green). B, Schematic representation of the case-control study design using WGS. Variants were identified using PEMapper/Caller and validated by GATK. Only shared variants between both pipelines were used. Following quality control measures of the raw WGS data, variant annotation was performed to identify rare (<1%) predicted LoF (loss of function), damaging splicing and damaging missense variants followed by filtering-based annotations on phastCons (conservation), CADD and CCRS (constrained coding regions) scores to identify MDRVs. Then weighted gene set based analysis and over representation analysis, ORA, was conducted. STRING analysis was performed to identify potential biological network interactions. C, Lateral side of mouse embryo at E10.5 with outline of tissues used for bulk RNA-sequencing (pharyngeal arches 2-6, PA2-6, blue; outflow tract and right ventricle, OFT+RV, green; left ventricle and atria, LV+atria, yellow).

### Ethics, consent and permissions

This study was approved by the Einstein Internal Review Board (Committee of Clinical Investigation, 1999-201) as well as each of the local institutional research ethics boards as part of the IBBC. We obtained de-identified DNA samples retrospectively collected with the informed consent of each of the subjects.

### Quality control of variants in raw WGS data

Details on the variant calling processes were provided previously (8). We performed a comprehensive QC analysis of the raw genotype data as described in Additional file 2: Figure S1. First, variants in the low copy repeat (LCR) regions and 3 million base pair 22q11.2 deletion region were removed. Single nucleotide variants (SNVs) were removed if genotype rate < 0.95 and Hardy-Weinberg equilibrium (HWE) < 10^−6^. Indels were removed if genotype rate < 0.97 and HWE < 10^−5^. Of note, we adopted a more stringent filtering criteria for indels as compared to SNVs because variant calling pipelines usually have a slightly lower confidence in identifying indels accurately. Monomorphic variants in the remaining 1,182 samples were then removed. A total of 21,695,115 of the 30,834,871 diploid variants in the raw data set passed QC procedures. The detailed composition of variants in the cleaned data are shown in Additional file 2: Figure S2. A total of 1,861,125 variants were identified as indels accounting for 8.62% of the total variation. Among them, 98.8% of the indels were within 10bp (Additional file 2: Figure S2A), which most variant calling pipelines have good sensitivity and specificity to make a call accurately (Fig. S2B). Most of the variants were rare with an alternative allele frequency (AAF) <=0.01 (Fig. 2C).

**Fig. 2.**
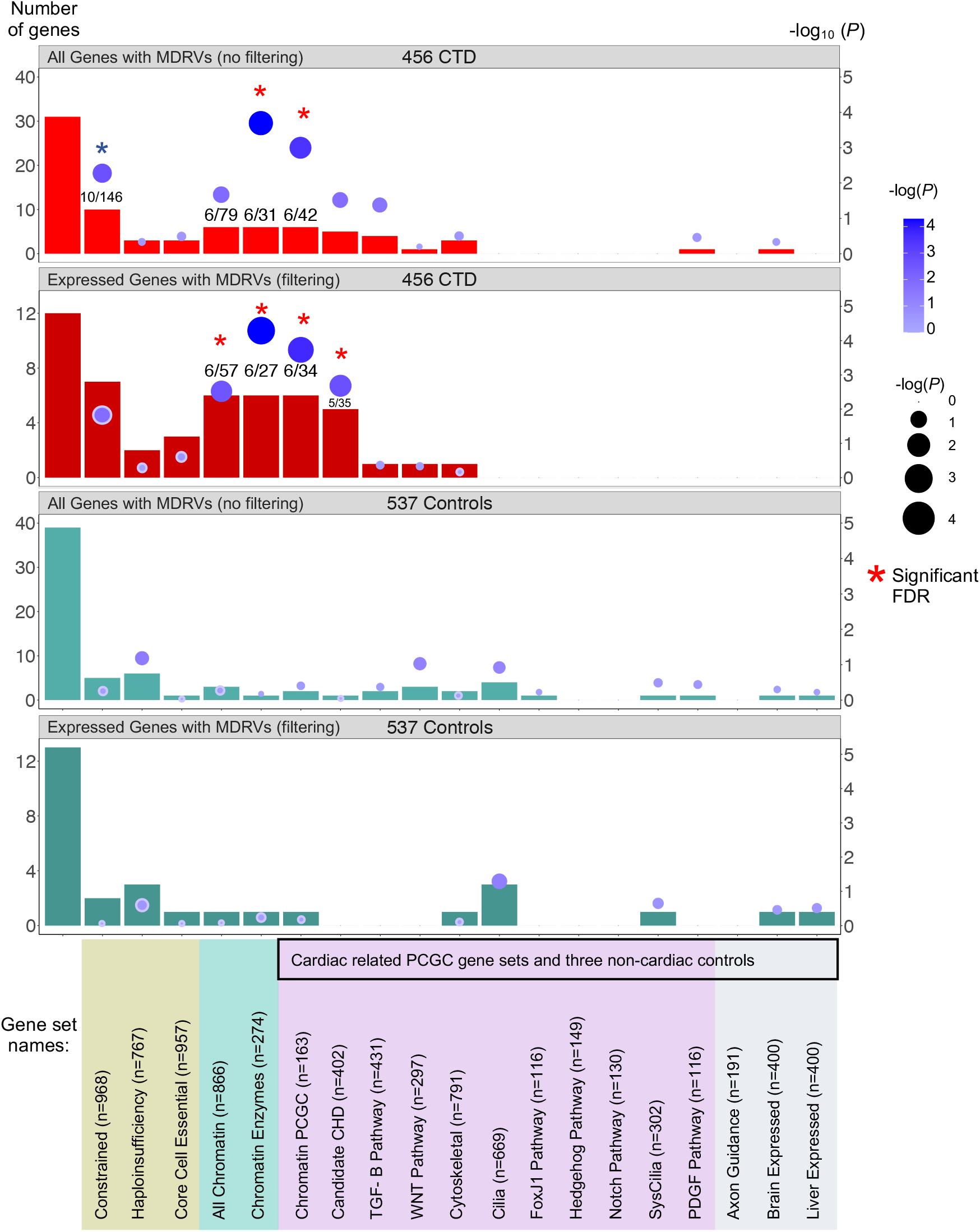
Over Representation Analysis (ORA) of recurrently affected genes identifies chromatin regulatory genes contributing risk to CTDs in 22q11.2DS. Three different sources of gene sets totaling 19 are indicated by color (gene sets used by the PCGC to investigate sporadic CHD^14^ are indicated by black box, as well as in lilac and in gray). The first bar on the left in each panel shows the total number of recurrently affected genes (n) among all affected genes (N). The rest of the bars indicate the number of recurrently affected genes within each gene set (k) versus the final number of affected genes with MDRVs (most damaging rare variants) for each gene set (M) as indicated (see Methods for more details). The top two bar graphs show ORA results without filtering by gene expression levels in CTD cases (red) and with filtering (dark red) followed by the same for controls (green and dark green, respectively). The numbers in some of the bars denote the number of recurrently affected genes contained within the specific gene set / the total number of affected genes in this gene set. The gene set analyses were corrected for multiple testing by false discovery rare (FDR). Red asterisks denote significance after FDR correction; blue asterisk denotes borderline significance (*P* = 0.057).

### Principal component analysis (PCA)

Details about PCA for this study have been described previously (8). A total of 879 (74.4%) subjects were Caucasian, 184 (15.6%) were of African descent or admixed and 119 (10.0%) were Hispanic (Additional file 2: Figure S3A). Further principal component analysis was stratified by case and control status (Additional file 2: Figure S3B) demonstrating that cases and controls are well matched by ancestry, indicating low probability of population stratification.

### Variant annotation and identification of the MDRVs

Protein truncating variant (PTVs) i.e., Loss of function (LoF) variants (14) including indel-frameshift, stop gain, splice donor, splice acceptor, stop loss, start loss were annotated in parallel by LOFFTEE(15), Bystro (16) and ANNOVAR (17). Damaging missense variants were annotated by ensemble score of MetaSVM (18). Damaging splicing variants were annotated by spliceAI (19) and two ensemble scores, ada (20) and random forest (rf) scores from the Ensemble database available from dbscSNV (21, 22). We are aware that the annotated functional variants may be enriched for false positives, for example, of sequence artifacts. Therefore, the putative functional variants were subjected to further filtering-based annotation to remove potential false positives and to obtain high-quality MDRVs. First, relatively less conserved variants were excluded at a phastCons score <0.5 and putatively less deleterious variants were removed with CADD (23) <10. Secondly constrained coding regions (CCRS) (24) restriction was further applied to prioritize and refine the MDRVs.

### *In silico* validation of the variants using GATK

Variants were called by GATK from the alignment to a Human Reference Genome (GRCh37/hg19) following default parameter settings in GATK release 4.0.0. To cross-compare variants from the two different pipelines, variants from PEMapper and PECaller were first lifted over from hg38 to hg19. A small proportion, 0.15% comprising 32,291 of 21,695,115 QC’d variants failed the liftover to hg19. The VCF files were split into individual VCF files for each sample, respectively. A total of 1,151 samples had variants/VCF files from both pipelines. Next, each pair of VCF files were cross-compared using Bcftools (https://samtools.github.io/bcftools/bcftools.html) as evaluated by concordance of chromosome, position, reference allele, alternate alleles and genotype i.e. in genotype mode or just the first four parameters (genotype mode =0). Indels were deemed as validated if >=10% of the base pairs overlapped.

### Gene level annotation

#### RNA-seq analysis of mouse embryonic cardiac development tissues

Genes contributing to CHD should be expressed in the embryonic pharyngeal apparatus, aortic arch and/or developing heart (25). We performed RNA-seq analysis using total RNA from the micro-dissected tissues of mouse embryos. Wild type embryos in a mixed SwissWebster/C57Bl/6 background at E10.5 (30-32 somite) were isolated in ice-cold Dulbecco’s phosphate-buffered saline (DPBS), then the pharyngeal arches 2-6 (PA2-6; cut along dorsa aorta), outflow tract together with the right ventricle (OFT+RV), and left ventricle plus atria (LV+Atria) of embryos were immediately micro-dissected and then were frozen separately and stored in the −80°C freezer. Each sample (each biological replicate) for RNA isolation was pooled from four embryos and three biological replicates were prepared for the RNA-seq. Total RNA was extracted using TRIzol and miRNeasy Mini Kits (QIAGEN, 217004), and on-column DNase I digestion (QIAGEN, 79254) was performed before eluting RNA in the RNase/DNase free water and all samples passed quality control measures. Library preparations and Illumina sequencing were performed at the Einstein Epigenomics Core Facility (https://einsteinmed.org/departments/genetics/resources/epigenomics-core.aspx). The libraries were prepared using the KAPA stranded RNA-seq Kit with RiboErase (HMR) (KAPA-Roche) following the protocol provided in the Kit. Ribosomal RNA was removed before library preparation. Quality of libraries was examined by Qbit (fluorometric quantitation; Invitrogen), bioanalyzer (Agilent 2100) and qPCR (Roche light cycler) and all passed quality control. Then the libraries were multiplexed and sequenced using the Illumina NextSeq 500 system as 2×150 bp paired ends. For RNA-seq analysis, the mean gene expression level from the three tissues was used to prioritize protein-coding genes, and the log transformations of the mean expression levels were used as weights for the weighted gene-set based test.

#### Gene constrained scores

Genes that are crucial for the development of an organism and survival will be depleted of LoF variants in natural populations, whereas non-essential genes will tolerate their accumulation (15). We used the latest gene-level constraint score, rank summation with re-sorting (VIRLOF) (26), which was based on deeply curated LoF variants from 125,748 exomes and 15,708 genomes of sequence data aggregated by gnomAD and the combination of LoF observed/expected upper bound fractions (LOEUF) (15) and gene variation intolerance rank (GeVIR) (26).

### Gene sets (19 in three clusters)

#### Three constrained score based gene sets

The Constrained Genes gene set was curated from the top 5^th^ percentile of VIRLOF gene level matrix (n= 968). These data were derived from population genetics approaches (15). The Essential Genes gene set included the shared core set of essential genes derived from three independent large-scale screens to assess the effect of single-gene mutations on cell viability or survival of haploid human cancer cell lines, cell-based essentiality (27–29) (n=956). The Haploinsufficiency Genes gene set was compiled based on the top 5^th^ percentile (n=767) from the genome-wide haploinsufficiency score (HIS) (30).

#### Two chromatin related gene sets

It has been established that *de novo* variants in chromatin related genes are associated with the pathogenesis of sporadic (25, 31) and syndromic CHD (32). We therefore downloaded all genes with chromatin function and modification related terms from GSEA (https://www.gsea-msigdb.org/gsea/index.jsp), totaling 866 chromatin related genes from 81 GO and REACTOME terms including chromatin (de)acetylation, (de)methylation, (de)phosphorylation, (de)ubiquitylation, chromatin remodeling, chromatin assembly or disassembly. We named this gene set as All Chromatin Regulatory Genes (n=866). We note that the small Chromatin Genes (n=163) gene set used by the PCGC comprises a small subset of the All Chromatin Regulatory Genes gene set. For the second gene set we focused on one Reactome term, Chromatin Modifying Enzymes (n=274), because it included many chromatin modifying enzymes. The first and larger gene set covers most chromatin genes and serves as a hypothesis-free gene set, while the second, represents a smaller subset of the first.

#### Eleven gene sets with recognized relevance to pathogenesis of sporadic CHD and three non-cardiac gene sets

From an original 19 gene sets previously curated and employed to investigate the genetic architecture of sporadic CHD (33), we selected 14, each having over 100 genes. These include 11 known pathways with recognized relevance to the pathogenesis of sporadic CHD, and three non-cardiac control gene sets. The cardiac gene sets include: Candidate CHD Genes (excluding cilia genes, n=402), Chromatin Genes (n=163), Cilia Genes (n= 669), SysCilia Genes (n=302), which is a subset of Cilia Genes, Notch Associated Genes (n=130), TGF-B Genes (n=431), Cytoskeletal Genes (excluding cilia, n=791), WNT Signaling Genes (n = 297), Hedgehog Signaling Genes (n=149), FoxJ1 Genes that consist of mobile cilia genes (n=116), Platelet Derived Growth Factor (PDGF) Signaling Genes (n=116). The three non-cardiac gene sets are Axon Guidance Genes (n=191), Brain Expressed Genes (n=400) and Liver Expressed Genes (n=400), serving as controls.

#### Mouse gene liftover to human orthologs

Mouse genes were lifted over to human orthologs using the R package, biomart (9, 34). For genes with either multiple mapping or missing mapping, web crawler was applied to automatically search for possible orthologs in NCBI genes (https://www.ncbi.nlm.nih.gov/gene).

#### Overrepresentation Analysis (ORA)

ORA is a widely used approach to determine whether known pathways or gene sets are over-represented (enriched) in an experimentally derived gene list (35). We adopted a stringent criterion to first select only genes identified to have >= one MDRV in two or more CTD cases and no MDRV in controls (n=0 in controls). A similar criterion was used to select recurrently affected genes in controls, i.e., genes identified to have >= one MDRV in two or more controls and no MDRV in CTD cases (n=0 in cases). As most of the affected genes have one identified MDRV, these criteria can account for potentially random background noise for false positive MDRVs in the whole 22q11.2DS cohort. These restricted subsets of recurrently affected genes in the CTD group, and in the control group, were then evaluated by ORA for enrichment in the 19 gene sets independently. The ORA can be calculated by hypergeometric distribution (36):

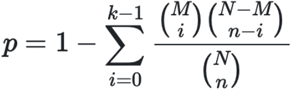

where N is the total number of genes included in the analysis as the background distribution; M is the final number of genes for each gene set among N, n is the number of recurrently affected genes; and k is the overlap of n and M, which is among the recurrently affected genes. For example, in the scenario of expressed genes in cardiac progenitor cells, in the CTD case group, N=413, M =27 for Chromatin Regulatory Enzymes (n=274), n=12 and k=6; while in the control group, N=466, M=30 for Chromatin Regulatory Enzymes (n=274), n=13 and k=1.

#### Weighted gene-based test

To reduce the noise/signal ratio, we employed a weighted gene-set based test to examine the aggregated burden of MDRVs in CTD cases as compared with controls. Specifically, let *Z* be *k* × 1 vector of z statistics of genes in a gene set obtained from the gene-based test for each gene, and *w* be *k* × 1 weighting vector of corresponding importance scores, more specifically, gene expression level in developing heart and related tissues that provide clues. Let 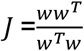 be a *k* × *k* importance score matrix. The statistic is defined as T = *Z*^T^*JZ*. For the 1,861 high-confidence MDRVs, we first performed a gene-based Fisher’s Exact Test between CTD cases and controls to obtain the gene-level *P* value and OR solely based on genetic data, then transformed this to the individual gene level statistic z. The log transformed mean expression level as described above was used as *w*, i.e., a given gene was assigned higher weight if it had a higher expression level in progenitor cells for cardiac development from RNA-seq analysis of E10.5 mouse embryos. An empirical gene set *P* value was calculated using 2,000 permutations by randomly shuffling case and control status. In each replicate, a T statistic was generated as described above for each gene set. The empirical *P* value for each gene set was calculated as *P* value = n / N, where n is the number of test-statistics as or more extreme than the observed test statistic and N is the total number of random permutations.

#### Multiple testing correction

We adopted the false discovery rate (FDR) to control for multiple testing burden. For the weighted gene set based test, we corrected for 19 *P* values. For the ORA, we adjusted for 19 × 4 =76 *P* values for the CTD and control group (×2), for filtered genes based upon expression and all genes (×2).

## Results

### Study design

To identify genetic modifiers, we analyzed whole genome sequence (WGS) data available for 456 cases with CTDs (22q11.2DS-CTDs) and 537 controls without CHD, all with 22q11.2DS (Fig. 1A). We applied PEMapper/PECaller (37), validated with GATK (38) (Fig. 1B), and then used public databases, comprehensive annotation software, and algorithms (Methods) to identify 1,861 high-confidence, predicted most damaging rare coding/splicing variants (MDRV, gnomAD alternate allele frequency (AAF) < 0.01; Fig. 1B; Additional file 2: Figure S4 and S5; Additional file 1: Table S1). We found 1.78 MDRVs per subject in 22q11.2DS-CTD cases and 1.85 MDRVs per subject in controls, with no significant overall excess of MDRVs in either group (*P* = 0.485, Additional file 2: Figure S6A). Male and female subjects had similar burden of MDRVs (1.81 versus 1.79, *P* = 0.803). Stratified analysis demonstrated there was no significant difference of MDRV burden between 22q11.2DS-CTD cases and controls within each of three main ancestry groups (all *P* >0.05, Additional file 1: Table S2), indicating that cases and controls were well matched for ancestry. The identified 1,861 MDRVs affected 1,261 genes, of which 83% had just one MDRV (1,048/1,261, Additional file 2: Figure S6B).

*Tbx1* is expressed in the pharyngeal apparatus (PA; arches 2-6) and in cells that migrate to the cardiac OFT and aortic arch arteries between mouse embryonic day (E)8-10.5 (39). To filter for genes most likely to be relevant to progenitor populations of the heart and *TBX1*, we generated RNA-seq data to determine the expression level of genes in the PA, OFT with right ventricle (OFT + RV), and/or left ventricle with atria (LV+atria) of mouse embryos at E10.5 (Fig. 1C). Of the protein coding genes in the mouse genome in each tissue, we considered the top 35% as expressed (with>=25 reads per million; Additional file 1: Table S3) and referred to them as cardiac progenitor cells.

### Chromatin regulatory genes identified with recurrent MDRVs in 22q11.2DS-CTD cases

We selected recurrently affected genes with MDRVs in at least two cases and no controls to help avoid possible random sequencing artifacts. A similar criterion was applied to the control group (two or more controls, no cases; Methods). In the 456, 22q11.2DS-CTD cases, there were 413 genes with MDRVs, 31 of which were recurrently affected. In 537 controls, there were 466 genes with MDRVs, and 38 were recurrently affected. We found 12 of 31 recurrent genes in cases (5.7% of cases), and 13 of 38 recurrent genes in controls, were expressed in mouse embryonic cardiac progenitors. Strikingly, half of the 12 recurrent genes with expression in cardiac progenitor cells in 22q11.2DS-CTD cases were chromatin regulatory genes: *NSD1, KAT6A, KMT2D, PHF21A, EP400* and *KMT2C*, of which some are associated with known syndromes that include CHD, supporting their candidacy (Table 1). Table 2 lists the detailed annotation of the MDRVs identified in the six recurrently affected chromatin regulatory genes, as well as the associated cardiac phenotypes. The most frequent cardiac phenotype was tetralogy of Fallot followed in frequency by right sided aortic arch (Table 2). Moreover, there were MDRVs in *KMT2C* in two additional 22q11.2DS participants (not included here as CTD cases or as controls), one with a ventricular septal defect (VSD) and one with an atrial septal defect (ASD; Tables 1 and 2).

**Table 1.** Twelve recurrently affected cardiac developmental expressed genes with MDRVs identified in CTD cases (and none in controls) in individuals with 22q11.2DS

|  |  |  |  |  |  |  |  |
| --- | --- | --- | --- | --- | --- | --- | --- |
| Gene | Full Name | Number of MDRVs | Genetic syndrome or disease | MIM ID | CHD present | Inheritance mode | CTD cases |
| **EP400** | E1A-binding protein, 400-kd | 1 |  |  |  |  | 2 |
| **KAT6A** | Lysine acetyltransferase 6A | 2 | Arboleda-Tham Syndrome | 601408 | yes | AD | 2 |
| **KMT2C** | Lysine-specific methyltransferase 2C | 2 | Kleefstra syndrome 2 | 606833 | yes | AD | 2 |
| **KMT2D** | Lysine-specific methyltransferase 2D | 2 | Kabuki Syndrome | 602113 | yes | AD | 2 |
| **NSD1** | Nuclear receptor-binding set domain protein 1 | 2 | Sotos Syndrome | 606681 | yes | AD | 2 |
| **PHF21A** | PHD finger protein 21A | 2 | Potocki-Shaffer syndrome | 608325 | occasional | AD | 2 |
| **CACNA1D** | Calcium channel, voltage-dependent, L type, alpha-1D subunit | 3 | Primary aldosteronism, seizures and neurologic abnormalities; PASNA | 615474 |  | AD | 3 |
| **CHERP** | Calcium homeostasis endoplasmic reticulum protein | 2 |  |  |  |  | 2 |
| **FBLN2** | Figulin 2 | 2 | Atrioventricular septal defect, susceptibility to; AVSD2 | 606217 | yes | Complex | 3 |
| **GTPBP4** | GTP-binding protein 4 | 1 |  |  |  |  | 2 |
| **RBMX** | RNA-binding motif protein, X chromosome | 1 | X-linked intellectual disability | 300199 |  | X-linked | 2 |
| **WASL** | WASP-like actin nucleation-promoting factor | 2 |  |  |  |  | 2 |
| non-duplicated |  | 24 |  |  |  |  | 26 |

**Table 2.**
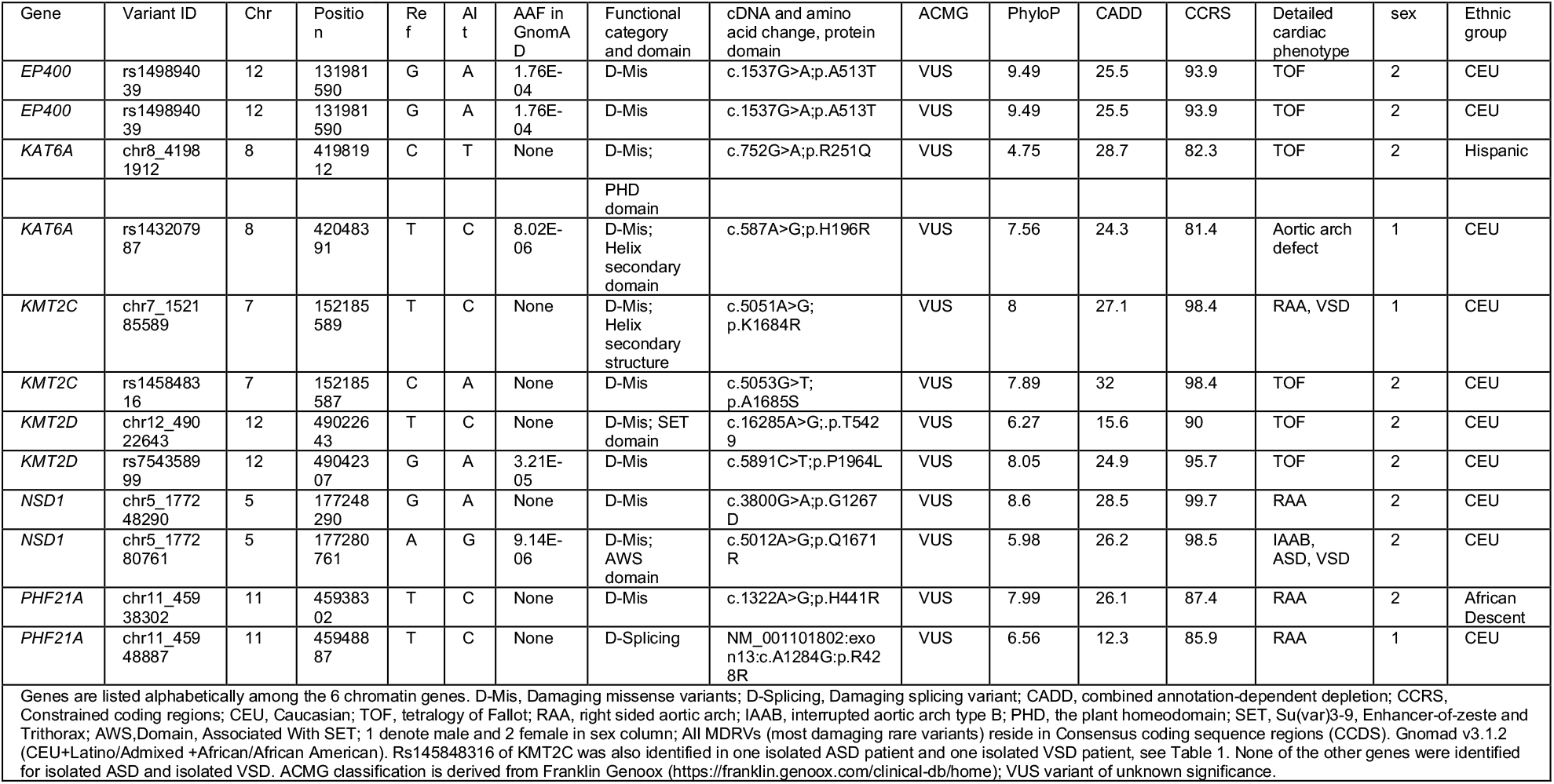
Identified MDRVs in the six recurrently affected chromatin genes in CTD cases

### Overrepresentation analysis of recurrent genes identifies chromatin regulatory genes in 22q11.2-CTD cases but not controls

Considering both the rarity of high-confidence MDRVs and relatively limited number of samples included, we next used a gene set based approach to test the aggregated effect of MRDVs in multiple functionally or conceptually connected genes in 19 gene sets, in cases and controls (Fig. 1B). The 19 gene sets were derived from three different sources of which had between 100 to 1,000 genes per set (Additional file 1: Table S4). This number of genes per set ensured sufficient power for our statistical tests. One source included constrained, haploinsufficiency, and essential gene sets that are under strong purifying selection (40), tend to play important roles in protein interaction networks (30, 41), and are highly enriched for disease genes (42). The second source was chromatin regulatory gene sets because some were identified in this study (Table 1). Additionally, *de novo* mutations in chromatin regulatory genes were found in individuals with sporadic CHD by the Pediatric Cardiac Genomics Consortium (PCGC) (25). One gene set consisted of 866 genes (All Chromatin Regulatory Genes, combined from several sources; Supplementary Table 4) and the other consisted of 274 genes and was a smaller subset (REACTOME term Chromatin Modifying Enzymes, Additional file 1: Tables S4 and S5). The third source comprised 14 gene sets (116 to 791 genes per set), derived from genetic studies of sporadic CHD by the PCGC, including genes found with mutations in one or more subjects (33). These 19 gene sets, though curated using differing concepts, are not mutually exclusive, as shown in the Venn plots in Additional file 2: Figure S7, indicating various degrees of overlap among different gene sets.

We then performed an overrepresentation analysis (ORA) of the recurrently affected genes on 19 gene sets (Fig. 2). We found that five of the six recurrent chromatin regulatory genes described above (Table 1; not *EP400*) contributed to findings for the Constrained Genes set with borderline significance (n=968; Fig. 2, panel 1, *P* =5.28×10^−3^). The chromatin gene sets showed significant overrepresentation, with all six genes shown in Table 1, either when filtered by expression in cardiac progenitor cells or not filtered (Fig. 2, panels 1 and 2; Additional file 1: Table S6). Further, four of these six genes contributed to findings for the Candidate CHD Genes set created by the PCGC (not *EP400* or *PHF21A*; n=402; Fig. 2, panel 2). None of the other gene sets were overrepresented nor were the three non-cardiac gene sets used as controls (Fig. 2; Additional file 1: Table S6). In the control 22q11.2DS samples, there was no enrichment of any gene set by ORA for the recurrently affected genes, regardless of developmental cardiac expression gene filtering (all *P* > 0.05 before correction; Fig. 2, panels 3 and 4), suggesting stochastic effects of MDRVs in other genes.

### Weighted gene set test expands chromatin regulatory genes identified in 22q11.2-CTD cases

We next employed a weighted gene set based test to evaluate whether high-confidence MDRV burden was significantly concentrated within any of the 19 gene sets. Again, we found chromatin regulatory genes contributing to 22q11.2DS-CTDs but identified a broader set of genes using this approach. The weighted gene set approach can reduce noise/signal ratio by prioritizing genes by their functional importance, and more specifically, by expression level during cardiac development, since genes in a gene set are not expected to contribute to disease equally (Methods). Genes affected with one or more MDRVs were included, thereby making it possible to expand the number of potential modifier genes. One gene set, Chromatin Regulatory Enzymes (274 genes), had significantly excess burden of MDRVs in 22q11.2DS-CTDs versus controls after false discovery rate (FDR) correction, with 24 identified MDRVs in cases and 14 MDRVs in controls (*P* = 5.00 × 10^−4^, Fig. 3, Additional file 1: Table S7); there was no significant enrichment of MDRVs in any of the three non-cardiac tissue gene sets (*P* >0.05 before FDR correction). The MDRVs that each affect one or more chromatin gene is either ultra-rare (n = 12, AAF <1.76×10^−4^) or novel (n = 30) in gnomAD (Additional file 1: Table S8). When taken together with ORA and weighted gene set approaches, we identified 42 MDRVs in 37 chromatin genes that occurred in 39, 22q11.2DS-CTD cases, accounting for 8.5% of 22q11.2DS-CTD cases in total. We found that 18 of the 37 chromatin genes are associated with known genetic syndromes that include CHD, supporting their candidacy (Additional file 1: Table S8).

**Fig. 3.**
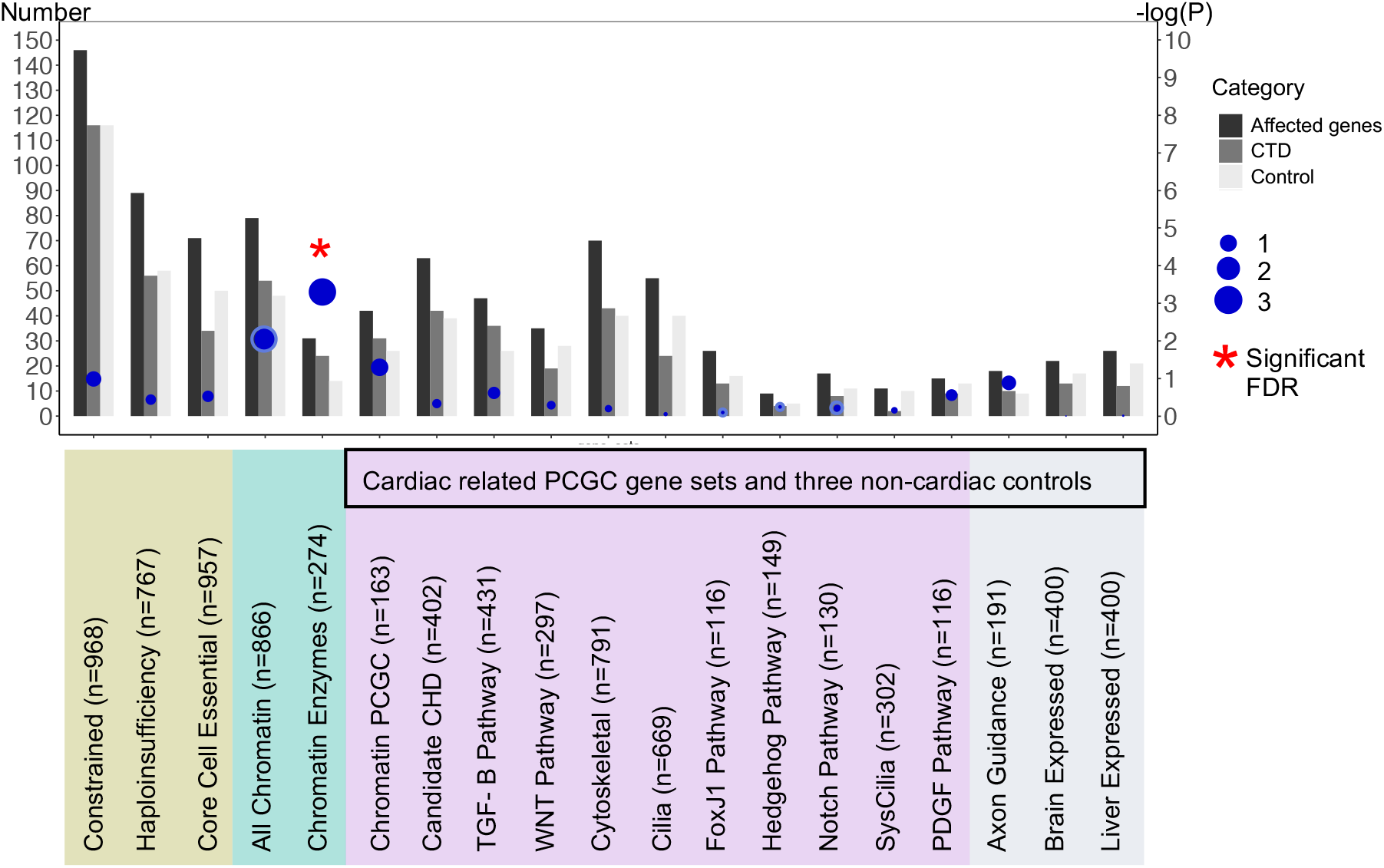
Identification of chromatin regulatory genes in 22q11.2DS by a weighted gene set approach. Three different sources of gene sets totaling 19 are indicated by color (gene sets used by the PCGC to investigate sporadic CHD are indicated; black box). Y-axis in the left denotes the number represented by the bars (three bars per gene set): the total number of genes included in each gene set for the weighted analysis, the total number of 22q11.2DS-CTD cases and controls. Genes were weighted by gene expression level (blue dots indicate *P* value; scale on Y-axis; red star is significant after FDR correction).

### Overlap of chromatin regulatory genes between 22q11.2DS-CTDs with sporadic CHD, but differences in pathogenicity of variants

We then compared the genes identified in 22q11.2DS-CTDs with chromatin regulatory genes found in cases of sporadic CHD from the PCGC. Sporadic CHD is defined by the PCGC as isolated cases in which neither parent is affected. Individuals with known syndromes, such as 22q11.2DS, were excluded, however subjects with isolated CHD or CHD that co-occurred with neurodevelopmental deficits or additional congenital anomalies were included (25). Most of the chromatin regulatory genes found in studies by the PCGC had *de novo* mutations and most occurred in sporadic CHD with neurodevelopmental disorders or extracardiac features (25). We cataloged genes that overlapped and found 13 chromatin regulatory genes in 22q11.2DS-CTD cases that were identified with *de novo* mutations in one or more subjects with sporadic CHD (90 chromatin regulatory genes among 2,871 CHD probands; Fig. 4A; Additional file 1: Table S9) (25, 31, 33). In an independent WES study of sporadic CHD to identify genetic risk factors, Sifrim and colleagues examined sequence variants in similarly categorized individuals with sporadic CHD (32). Nine genes in 22q11.2DS-CTD cases were found among 65 genes identified with *de novo* mutations in CHD with neurodevelopmental disorders or extracardiac features (32) (610 subjects, syndromic-CHD; S-CHD cases; Fig. 4A). In an integrated analysis(32), four chromatin genes in 22q11.2DS-CTD cases were identified among 16 found (16 *de novo* and inherited variants were found at the highest tier of significance among 1,891 probands with/out neurodevelopmental disorders or extracardiac features in S-CHD vs nonsyndromic, NS-CHD; Fig. 4A). When taken together, 14 genes were shared among the 22q11.2DS-CTDs, PCGC and Sifrim et al. studies (*CHD7, KAT6A, KMT2C, KMT2D, NSD1, SMAD4, TRRAP, EP400, IPO9, BRPF3, DNMT3A, HLTF, KDM4B, KMT2E*, Additional file 1: Table S9), thereby providing further support of their role in cardiac development and disease. This finding is particularly compelling when considering that the identification of rare variants was performed using different patient cohorts (22q11.2DS-CTDs, sporadic CHD in general population by the PCGC (25, 31, 33) and by Sifrim et al (32)), study design (case-control, trios, and both trios and singletons, respectively), variant types (unknown inheritance for 22q11.2DS-CTDs, *de novo* and inherited recessive variants for sporadic CHD studies) and variant annotation pipelines/statistical methods. Further, of the 14 genes among the three studies, eight of ten that are causative of genetic syndromes have CHD as a notable feature (43–48).

**Fig. 4.**
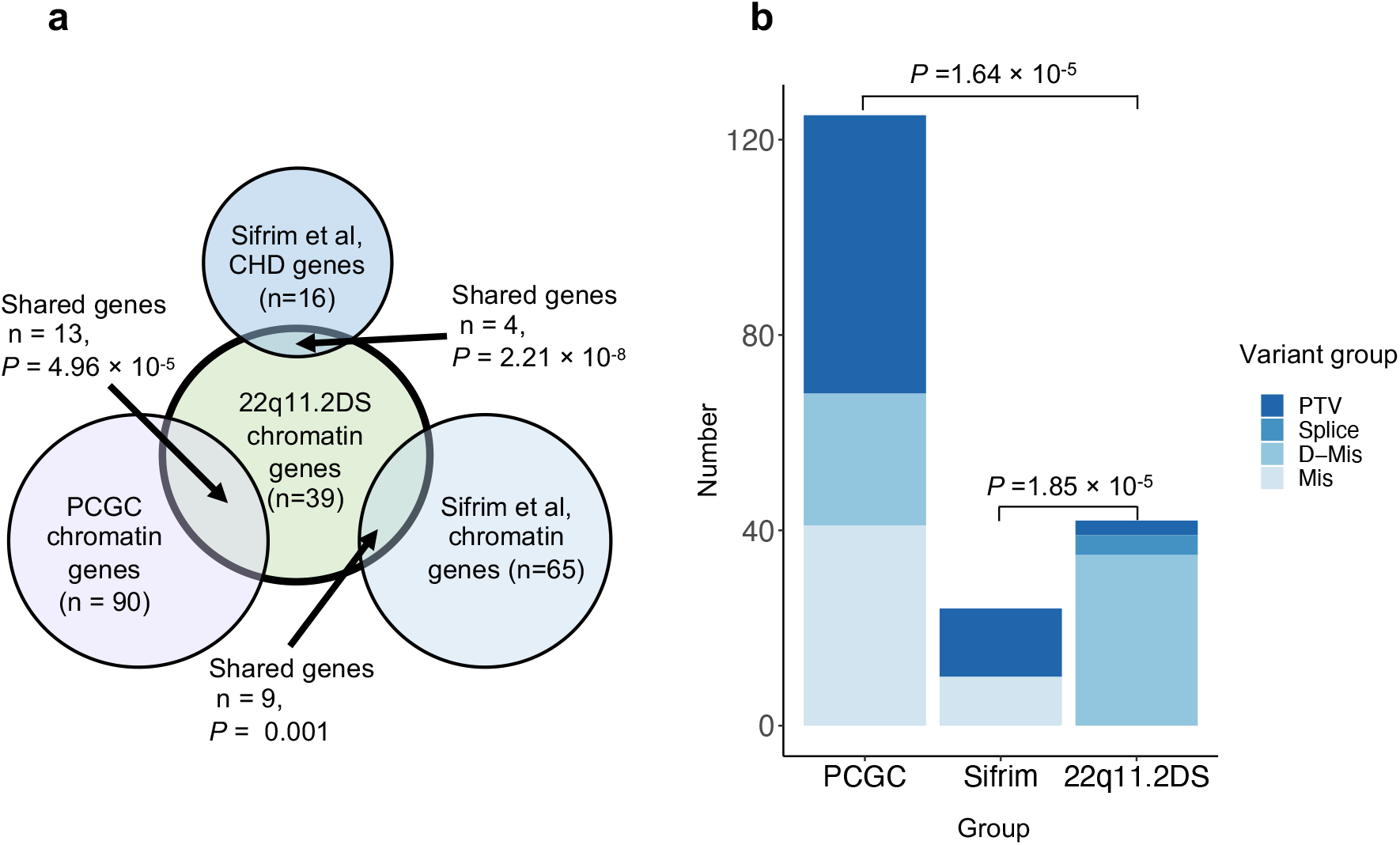
Chromatin regulatory genes shared with sporadic CHD in the general population. A, Venn plot showing the number of chromatin genes, as well as the number and *P* value for the overlap (arrow) of chromatin genes identified between 22q11.2DS-CTDs (green) and studies of sporadic CHD (PCGC-lilac and Sifrim-dark blue is from integrated analysis, light blue is *de novo* mutations in S-CHD). A total of 1,861 variants were found in 1,261 genes serving as the background of the analysis. B, Types of variants in chromatin genes (PTV is protein truncating, D-mis are damaging missense variants, mis is missense). A total of 57 PTVs were identified in 90 chromatin genes in sporadic CHD by the PCGC. In total, 14 PTVs were identified among 24 *de novo* variants in nine genes by Sifrim et al. A total of three PTVs were found among 42 variants in 22q11.2DS-CTDs. *P* values derive from two-Proportions Z-Test.

We next wanted to compare variants with pathogenicity identified in the clinical literature. Most of the MDRVs in 22q11.2DS-CTD cases were missense changes predicted to be damaging (Fig. 4B). From examination of clinical databases, 39 of 42 were considered variants of unknown significance (VUS; Table 2 and Additional file 1: Table S8). We then wanted to ask whether the types of variants in chromatin regulatory genes found in 22q11.2-CTDs might be similar to those found in studies of sporadic CHD. In contrast to 22q11.2-CTDs, protein truncating variants (PTVs; loss of function, LoF) were found more frequently in sporadic CHD by the PCGC (Fig. 4B) (25, 31, 49). This was similar for studies by Sifrim et al (32) (Fig. 4B). Further, many of the variants identified in studies of sporadic CHD, in particular for chromatin regulatory genes, were *de novo* heterozygous mutations, but the variants we identified were of unknown inheritance. This suggests that variants in chromatin regulatory genes in 22q11.2DS-CTDs may affect gene function but are not disease causing on their own, thereby serving as modifiers.

### Biological network connections between chromatin regulatory genes and *TBX1*

To understand the biology of the genes, and with respect to *TBX1*, we generated a functional protein network using STRING software (https://string-db.org: Fig. 5A). The network consists of the 37 chromatin regulatory genes we identified plus two additional genes, *CHD7* and *ATAD2B*, where we found MDRVs affecting more cases than controls in this study (two versus one, three versus two). As expected, there is an appreciable amount of functional coherence of these genes (*P* = 5.03×10^−14^). These chromatin regulatory genes have functions as histone lysine acetyltransferases and histone methyltransferases (Fig. 5B). We also identified sequence specific DNA binding proteins, suggesting cohesive but varied mechanisms by which these genes may increase risk for 22q11.2DS-CTDs (Fig. 5A and B).

**Fig. 5.**
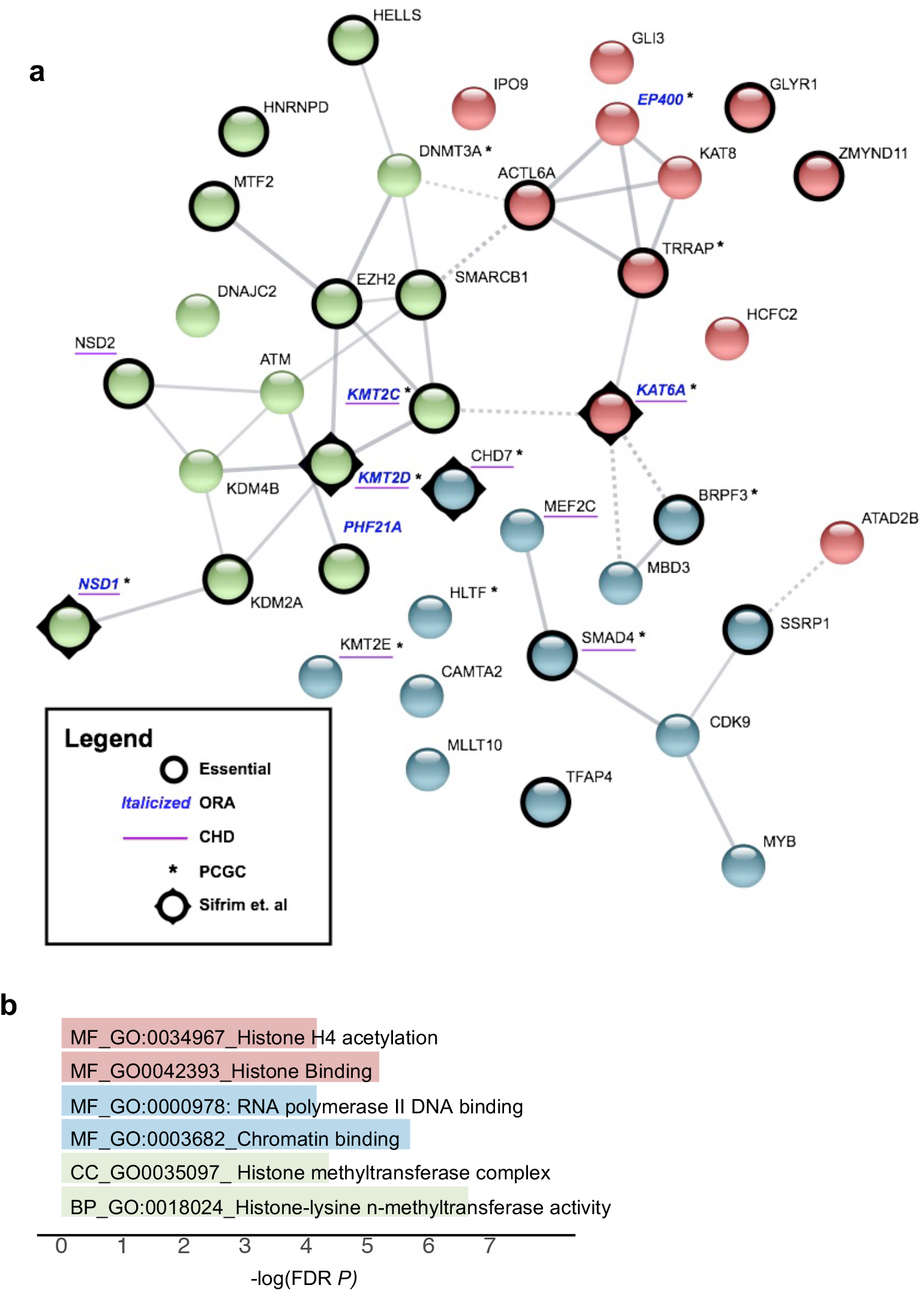
Chromatin gene network of modifiers for CTDs in 22q11.2DS. A, STRING image of 39 chromatin genes as identified in 22q11.2DS subjects with CTDs. Edges indicate both functional and physical protein interactions (Protein-protein enrichment p-value, 5.03×10^−14^). The types of interaction evidence for the network edges are indicated by the line color (text mining, experiments, database, co-expression, neighborhood, gene fusion and co-occurrence). The high confidence score of 0.700 was used to create the network. Kmeans clustering was used to generate three clusters (cluster 1, red; cluster 2, blue; cluster 3, green). The nodes based on confidence; with line thickness indicates the strength of the support of the data in STRING. Six genes found by ORA are indicated (Italic blue font for gene names), chromatin genes found with *de novo* mutations in one or more cases with sporadic CHD (with asterisk next to the gene names). Constrained genes are indicated (Black circle in nodes) and candidate sporadic CHD genes are shown (underscore for the gene names). B, Representative most significant gene ontology terms from the STRING image, color coordinated according to the STRING image (molecular function, MF; cellular component, CC; biological process, BP). FDR, false discovery rate, −log10 *P* value.

A total of 22 of the chromatin regulatory genes and five of the six recurrent genes were among the most Constrained Genes gene set, significant overlaps for these shared genes (overlap *P* = 1.27 ×10^−17^ and 6.27 ×10^−6^, respectively; Fig. 5A). Additionally, nine are implicated in causing sporadic CHD from other studies in the literature (25, 31, 50) (overlap *P* = 7.05 ×10^−5^; Fig. 5A). Further, we show the overlap between the genes for sporadic CHD from the PCGC and the four from Sifrim et al (Fig. 5A; Additional file 1: Table S9).

We next examined expression levels of the chromatin genes found in 22q11.2DS-CTDs in cardiac progenitor tissues. *TBX1* is expressed in the progenitor cells of the pharyngeal apparatus (PA) that migrate to the cardiac OFT during early embryogenesis (2). A logical hypothesis based upon results of this study is that altered chromatin regulatory genes in 22q11.2DS-CTDs may modify expression of *TBX1* and/or downstream genes, disrupting normal cardiac development. If this is the case, such chromatin modifiers should be expressed in the same cell types as *TBX1* during the same developmental period. Most chromatin regulatory enzymes are widely expressed but they may show enrichment in certain cell types or tissues. We therefore examined expression in the PA, OFT + RV and LV + atria. We found 36 of the 39 genes show enriched expression in *TBX1* relevant tissues, specifically the PA as compared to either the OFT + RV or the LV + atria, where *TBX1* is not expressed (Fig. 6; Additional file 1: Table S3) providing support for our hypothesis that chromatin regulatory genes might act in the genetic and epigenetic pathways of *TBX1* function.

**Fig. 6.**
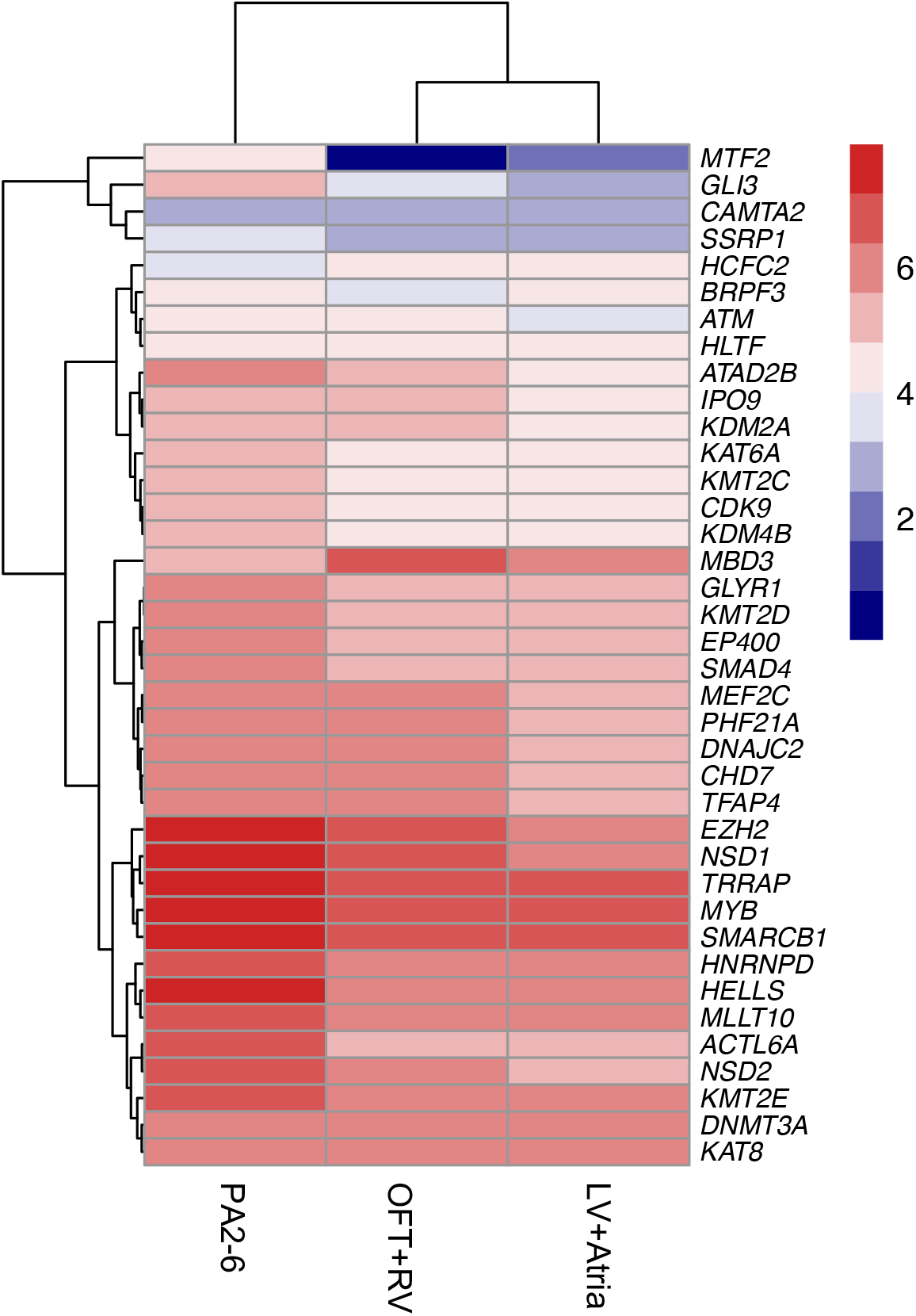
Expression of chromatin regulatory genes is enhanced in cardiac progenitor cells of the pharyngeal apparatus. Heatmap plot of RNA-seq results of genes from Figure 5 by the log 2 transformed expression level in PA2-6, OFT+RV and LV+atria as indicated with highest red to lowest blue color. One gene, *HNRNPD* is significantly highly expressed than the rest of genes KPM= 507, 334 and 296, respectively, as the rest expression KPM value ranging from 1-197.

## Discussion

This is the largest study to date in which WGS was used to identify genetic modifiers of 22q11.2DS-CTDs. From this study, we uncovered rare potentially damaging variants (MDRVs) in chromatin regulatory genes that we suggest serve as modifiers in 8.5% of individuals with 22q11.2DS-CTDs. Some of the chromatin regulatory genes identified are co-expressed in the same cell type with *TBX1*, encoding a transcription factor and the main gene for 22q11.2DS. We generated a model to explain the shared types of chromatin regulatory genes and possible mechanisms by which some of these genes may interact with TBX1 (Fig. 7).

**Fig. 7.**
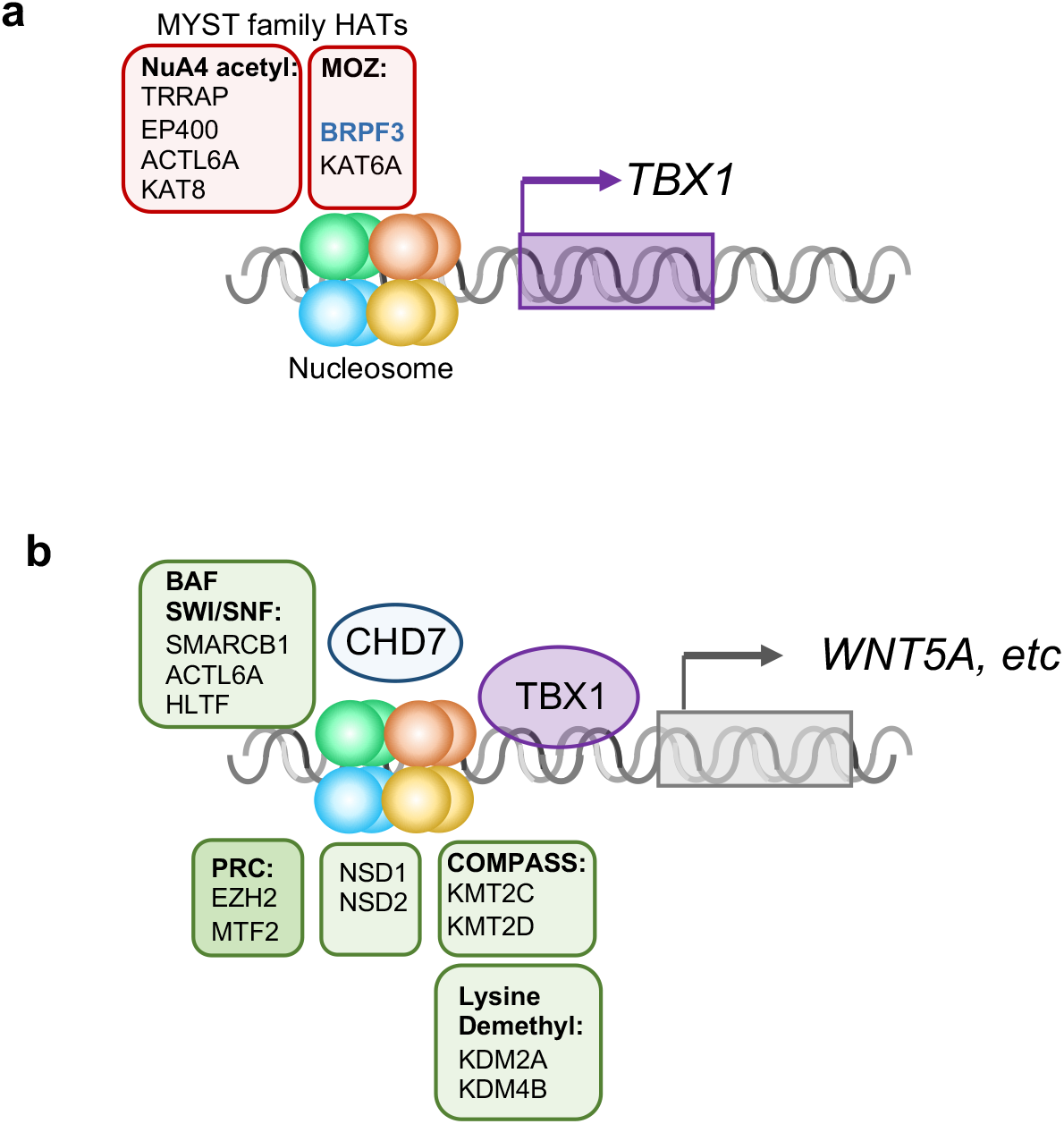
Model of chromatin regulators that mediate TBX1 function. Protein complexes involved in histone modifications (NuA4 histone acetyltransferase complex-NuA4 acetyl; Monocyte leukemia zinc finger complex-MOZ; BRG1/BRM complex (SWI/SNF), BAF; Polycomb repressive complex, PRC; Complex of proteins associated with Set1, COMPASS) are shown (similar colors as Figure 5A-STRING image) surrounding a representative nucleosome. A, MYST family proteins involved in histone acetylation with respect to regulation of *TBX1* expression. B, TBX1 protein regulates expression of downstream genes including WNT5a, via chromatin regulators that belong to several classes as indicated. This is mediated in part, by physical interaction with CHD7. DNA is shown as a gray double helix. Gene activation is shown as an arrow and repression as a cross bar.

We identified two main classes of chromatin regulatory proteins with MDRVs as illustrated in Fig. 7. Fig. 7A, shows the histone acetyltransferases (HATs) that harbored MDRVs in this study. These HATs all share a MYST domain and have important roles in transcriptional activation (51). We identified MDRVs in genes in two MYST different histone acetyltransferase families, the MOZ and NuA4 complexes (Fig. 7A). One of the genes in the MOZ complex is *KAT6A*. Of interest, it was found that *Kat6a* genetically interacts with *Tbx1* in formation of the aortic arch in mouse models and the mechanism is by regulation H3K9 acetylation in the *Tbx1* locus resulting in activation of gene expression (52) (Fig. 7A). It is therefore possible that MDRVs in these HATs mediate transcriptional activation of *TBX1* and/or downstream genes.

In Fig. 7B, we show the second main class of chromatin regulatory genes. We included lysine methyltransferases that we identified with MDRVs and a possible role with TBX1 protein in regulating transcription of downstream genes. KMT2C and KMT2D are two methyltransferases that we identified with recurrent MDRVs (Fig. 7B). Heterozygous gene inactivating mutations in *KMT2D* cause Kabuki syndrome in humans. KMT2C and KMT2D are part of COMPASS-Like family of H3K4 histone lysine methyltransferases (53). It was shown that TBX1 physically interacts with KMT2C to mediate H3K4me1 methylation to activate downstream genes such as *WNT5A* (54) (Fig. 7B). The H3k4me1 mark is for poised enhancers that are further modified to promote gene expression (55). Other proteins we identified with MDRVs are members of the PRC (Polycomb) complex that represses transcription, the BAF complex that alters nucleosomes (SWI/SNF) (56) and NSD1 and NSD2, which are involved in lysine methylation (57). EZH2 is a member of the polycomb complex that methylates H3K27 and *Ezh2* genetically interacts with *Tbx1* for mouse cardiovascular development (58), supporting a connection to *Tbx1. CHD7* is the gene for CHARGE syndrome and functions as an ATP-dependent chromatin remodeler that interacts with the BAF complex (56) as shown in Fig. 7B. TBX1 genetically and physically interacts with CHD7 for formation of the aortic arch (59). Of interest, *KAT6A* and *CHD7* were also among the 26 genes identified for tetralogy of Fallot patients based on reanalysis of whole exome sequence data from 811 subjects (60). Therefore, some of the genes we identified have been shown to be in the molecular pathway of TBX1, supporting their candidacy as modifiers.

*De novo* gene inactivating mutations in chromatin regulatory genes occurred in 3% of individuals in previous sporadic CHD studies from the PCGC (25, 31, 49). Interestingly, we found an even greater occurrence of variants in chromatin regulatory genes in our 22q11.2DS cohort, with MDRVs identified in 8.5% of the 22q11.2DS-CTD cases. We previously identified chromatin regulatory genes from whole exome sequence from 89 individuals with 22q11.2DS and tetralogy of Fallot versus controls, although there were differences in the genes themselves (61). Half of the genes that overlapped between our study, reported here, of 22q11.2DS-CTDs and studies of sporadic CHD from the PCGC, are causative of known syndromes in which CHD is an associated feature (43–48). For some of the genes we identified in 22q11.2DS-CTDs, there were between 5-17 CHD cases with mutations among 2,391 sporadic CHD trios (*CHD7* [n=15], *HLTF* [n=5], *KMT2D* [n=17] and *NSD1* [n=6]), but for others were one to three subjects affected (*DNMT3A* [n=1], *BRPF3* [n=1], *EP400* [n=1], *KAT6A* [n=2], *KMT2C* [n=3], *KMT2E* [n=1], *SMAD4* [n=1] and *TRRAP* [n=1]) (33). Our study of 22q11.2DS-CTDs therefore provides more support for some of the chromatin regulatory genes and their contribution to the etiology of CHD. The results in this study and previous work (10, 61), supports the idea that genetic modifiers for 22q11.2DS-CTDs are inherited as a complex trait, similar to risk factors for sporadic CHD. However, in our study most of the variants were missense changes in chromatin regulatory genes that are predicted to be damaging, while for sporadic CHD, most were protein truncating gene inactivating mutations (31, 33, 50).

In addition to *de novo* heterozygous mutations in chromatin regulatory genes, recessive inherited cilia genes were identified in the PCGC study by taking gene set approaches (31, 33). Some of the subjects with recessive mutations in cilia genes had laterality defects such as heterotaxy, while others had CTDs (33). We did not find evidence for heterozygous MDRVs in cilia genes using the same cilia gene sets as previously analyzed (33) and this suggests that 22q11.2DS-CTD modifiers do not parallel all the different types of genetic risk factors for sporadic CHD.

One of the limitations of this study is that we have a relatively small sample size which required us to test the aggregated effect MDRVs at the gene set level. However, by examining 19 gene sets, we found significant enrichment of MDRVs burden in chromatin regulating genes, which suggests it is not the overall burden but rather the regional MDRV burden of genes which plays critical role in TBX1 regulation network as supported by the transcriptomic profiling data, that makes the cardiac phenotypic difference in the 22q11.2DS cohorts. The 19 gene sets in this study not only served as the testing unit to increase statistical power for the aggregated effect of mutations but also as gene level annotation, by which we found the chromatin regulatory genes identified for 22q11.2DS-CTDs also had appreciable constrained features, implicating their intolerance to variation. Furthermore, we didn’t find enrichment among the gene sets in the 22q11.2DS control group.

In summary, our findings suggest that disturbance of a *TBX1* gene network by the presence of MDRVs in chromatin regulatory genes may disrupt cardiac development in a significant subset of individuals with 22q11.2DS, thereby serving as genetic modifiers. Since some of these chromatin genes were found in only one or a few individuals with sporadic CHD, our work provides more support for their candidacy as disease genes beyond many of their known roles in established genetic syndromes. This study therefore emphasizes the shared mechanisms involving the *TBX1* gene network in CHD.

## Code availability

The core computational pipelines used for quality control, annotation, and statistical analysis of the WGS data was coded in python 2.7. The weighted gene-set based analysis and plot of all figures were done in RStudio. Both the python code and R code are available upon request.

## Data availability

Data supporting the findings of this work are provided in Supplementary Data Tables. The raw WGS data for the 22q11.2DS cohort was previously deposited on NIMH Data Archive (https://nda.nih.gov/study.html?id=938).

## Acknowledgements

We acknowledge support from the Genomics and Epigenomics Core facilities at Einstein. This work was supported by a Leducq Foundation grant and NIH grants P01HD070454, R01HL157157, R01GM125757 and U01MH101720. National Institute for Mental Health (MBMvdB, 5U01MH101724). D.H.S. has been funded by Instituto de Salud Carlos III through the project “PI18/00847” (cofunded by the European Regional Development Fund/European Social Fund’s “A way to make Europe”/” Investing in your future”). JASV is supported by NIMH grant U01MH119741-01 and SickKids Psychiatry Associates Chair in Developmental Psychopathology. JB is supported by a senior investigator fellowship of FWO Flanders. MJO was supported by Medical Research Council UK, Centre Grant No. MR/L010305/1, Program Grant No. MR/P005748/1. We thank The Centre for Applied Genomics (Toronto, Canada) for informatics expertise. We thank Xianhong Xie for statistical advice. Steven T. Warren was a co-PI of the 22q Sequencing Consortium and significantly contributed to this work prior to his untimely death in June of 2021. We thank Gloria Stoyanova for the help with generating the STRING figure.

## Extended IBBC author list

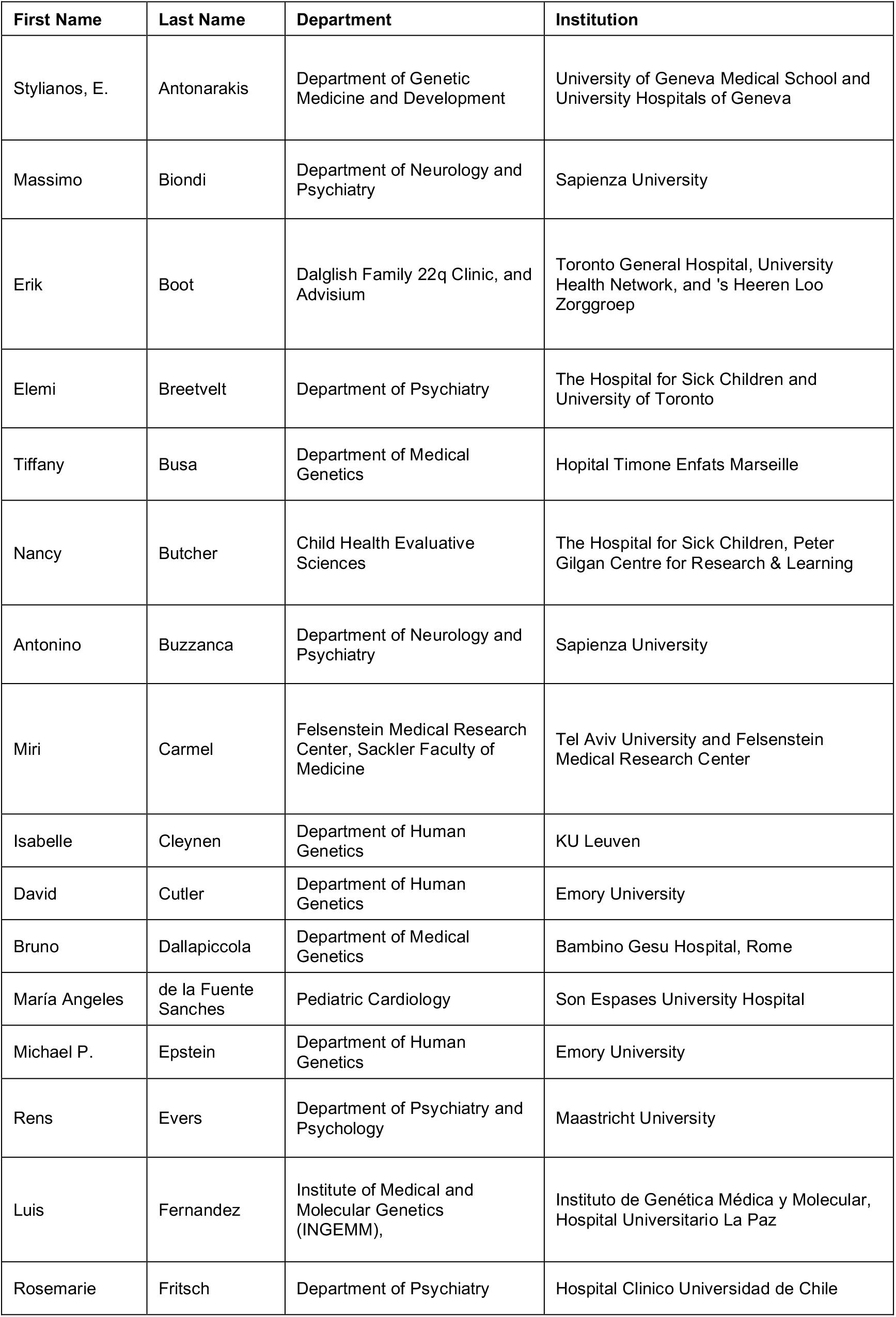
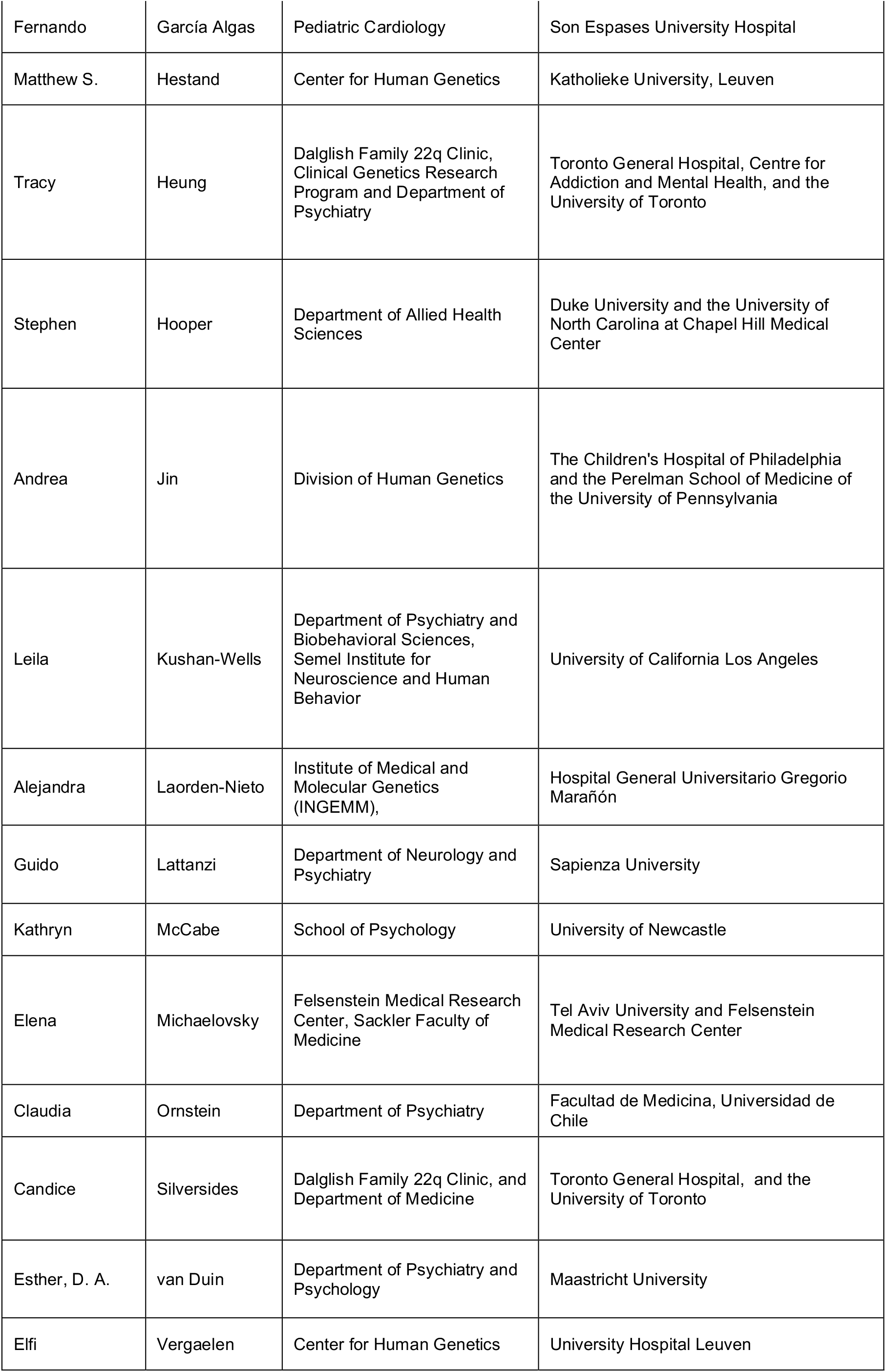
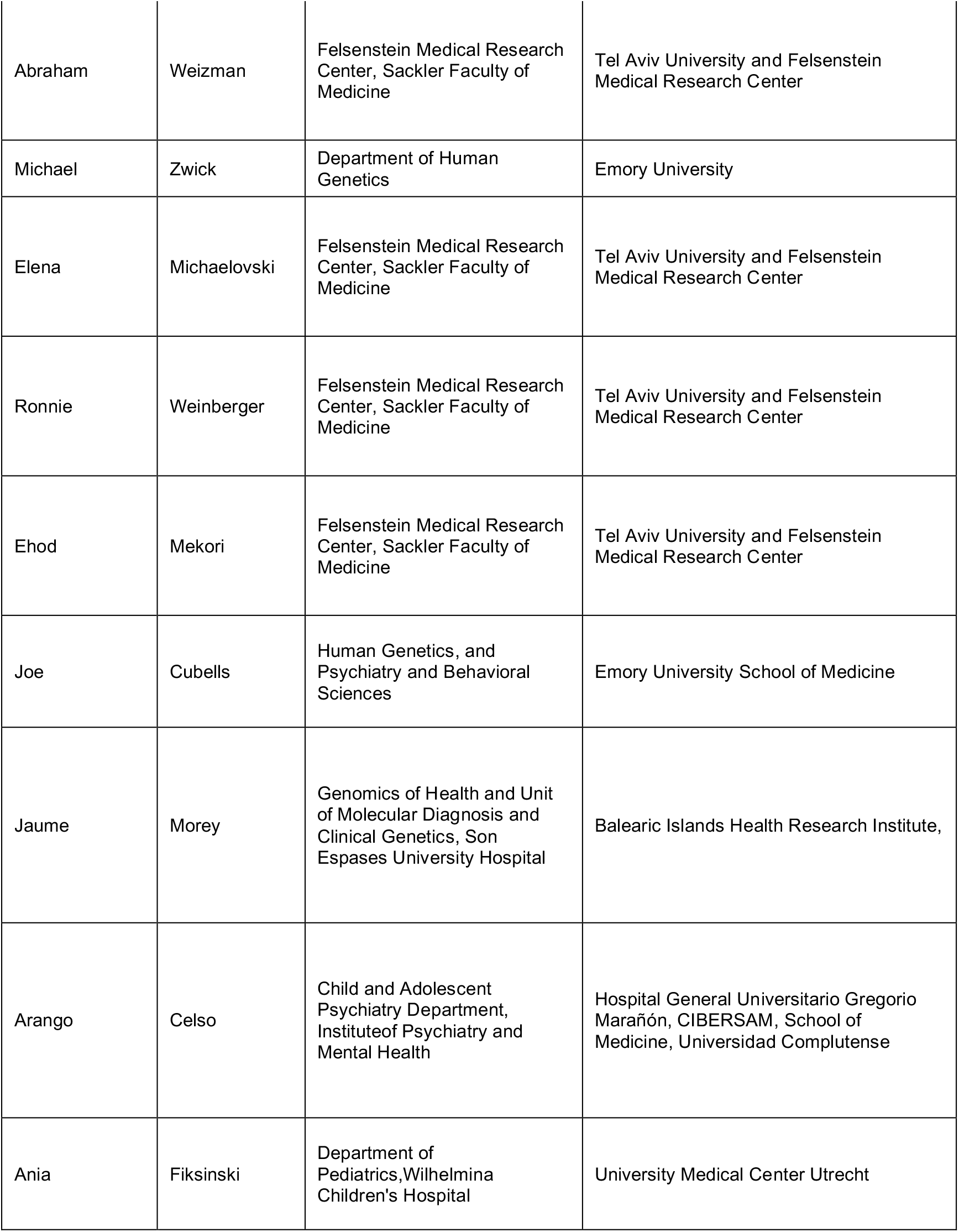

## Author contributions

BM conceived the idea. BM and YZ designed the study. YZ developed the methodology and set up the analytical framework and drafted the manuscript. YZ and YW programmed and implemented the whole analysis. LS generated and analyzed the mouse RNA-seq data. TW provided statistic consultation. HRJ called the variants in PEMapper/PECaller pipeline. CM called the variants in GATK pipeline. DZ provided analysis of the RNA gene expression data. The remaining of the authors identified, recruited and obtained study participant samples from multiple collaborating sites, provided phenotypic data and helped generate genomic raw data. All authors contributed to subsequent version and have read and approved the manuscript.

