## Supplemental Figures for "Chromatin regulators in the TBX1 network confer risk for conotruncal heart defects in 22q11.2DS and sporadic congenital heart disease"

### Figure S1

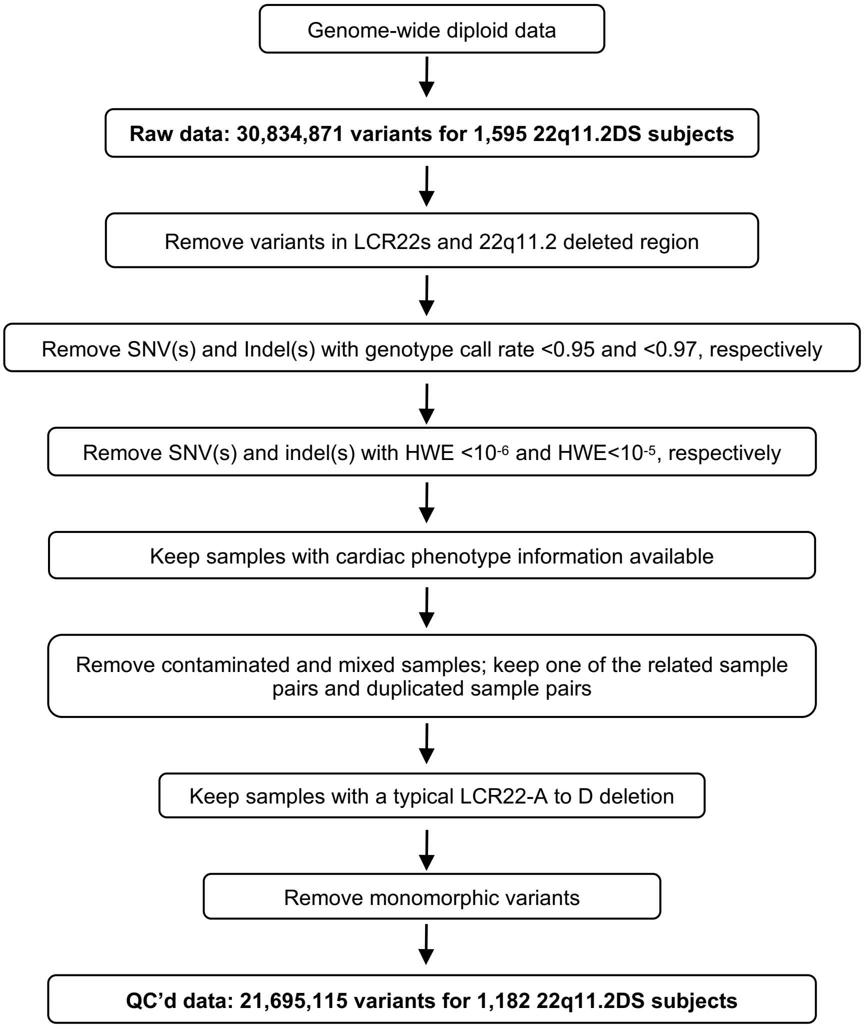


**Figure S1. Detailed quality control (QC) procedures applied to unprocessed WGS data.** Flow diagram of QC procedures used in this study. LCR22: Low Copy Repeats on chromosome 22q11.2; LCR22-A and LCR22-D mark two LCR22s that are three million base pairs apart, in which flank the most typical deletion. HWE: Hardy-Weinberg equilibrium; QC: quality control.

Figure S2

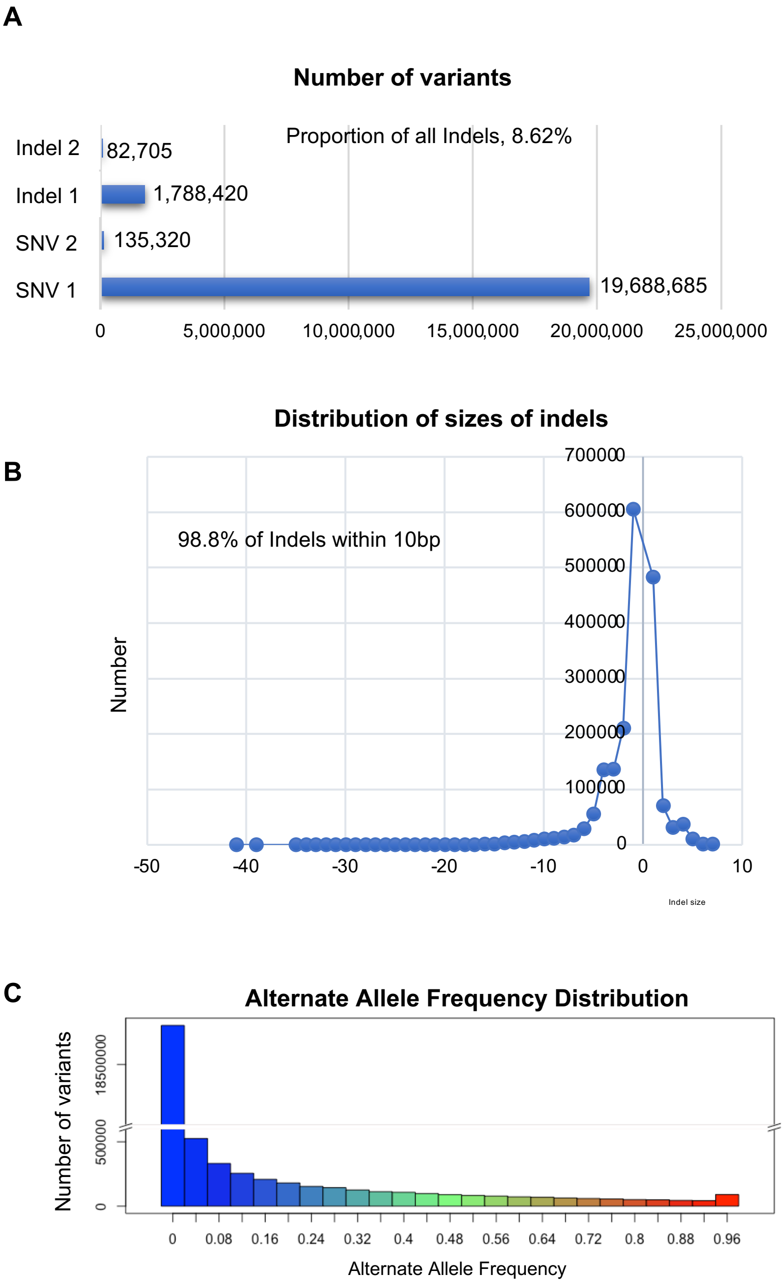


**Figure S2. Summary of quality controlled WGS data.** A, Bar plot shows the total number of the four variant categories, biallelic SNV (single nucleotide variant) and SNVs with more than one alternate allele; biallelic indel and indels with more than one alternate allele. B, Distribution of indels based on size. A total of 98.8% of all indels are within 10bp. C, Histogram shows the distribution of the alternative allele frequency (AAF) and the vast majority are rare variants (AAF<0.01).

### Figure S3

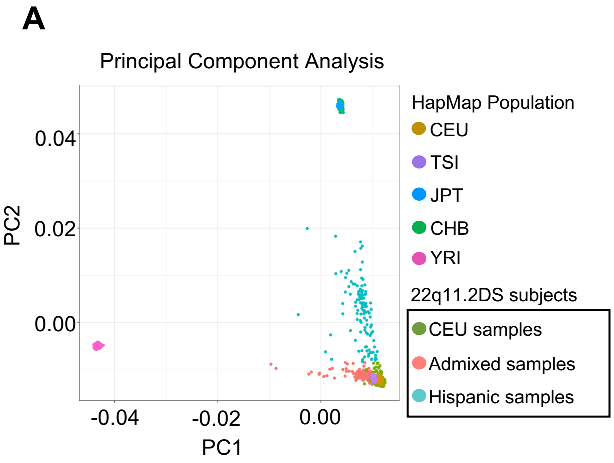


| 22q11.2DS on PCA | Subjects (%) |
|---|---|
| Caucasian | 879 (74.4%) |
| Admixed | 184 (15.6%) |
| Hispanic | 119 (10.0%) |

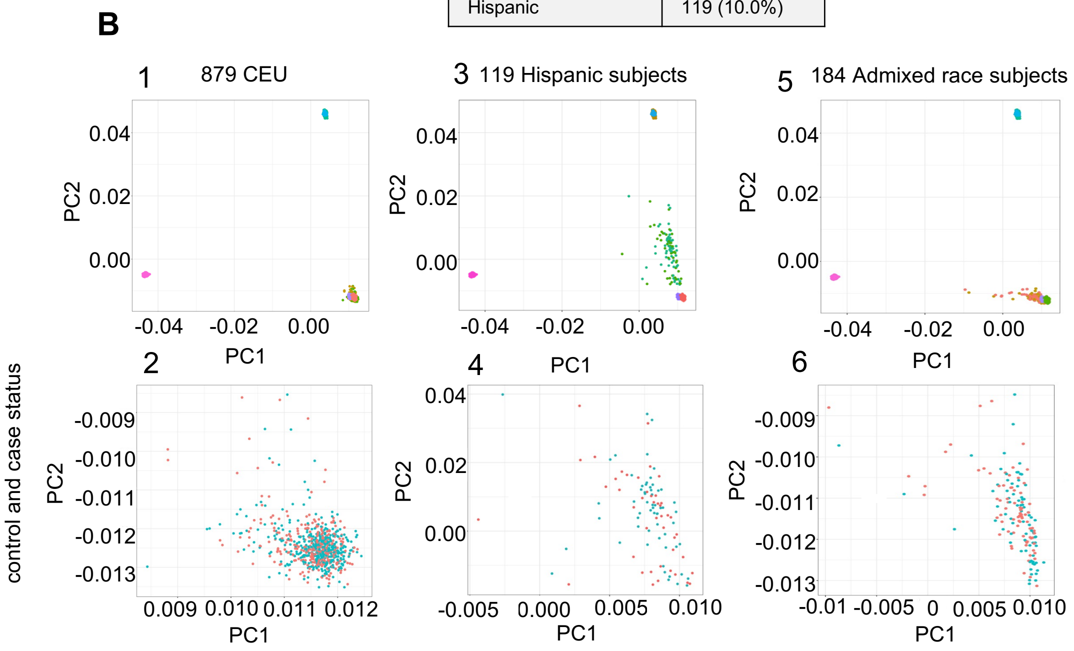


**Figure S3. CTD cases and controls are matched for each ancestry group as determined by principal component analysis (PCA).** A, PCA of 1,182 subjects to identify composition of race and ancestry. Reference population data is from HapMap Phase III and the 22q11.2DS cohort (22q11) data is indicated within CEU (Caucasian), AD (African descent), admixed samples and Hispanic samples based on the top PCs (black box). Table illustrates numbers**. B,** PCA within each ethnic group: B1, Scatter plot of 879 CEU samples by case control status (gold and green) against the same sets of HapMap samples plotted in a**,** B2, Zoomed in scatter plot of 879 CEU samples by case and control status (red and turquoise). B3, Scatter plot of 191 Hispanic samples by case control status (light green and green) against the same sets of HapMap samples plotted in Fig. 1b, B4, Zoomed in scatter plot of 191 Hispanic samples by case and control status (red and turquoise). B5, Scatter plot of 184 AA and mixed ancestry samples by case control status (light green and green) against as the same set of HapMap samples plotted in Figure 1B. B6, Zoomed in scatter plot of 184 AD and mixed ancestry samples by case and control status (red and turquoise). AD: African Descent; CEU: Utah residents with ancestry from Northern and Western Europe; TSI: Tuscan in Italy; YRI: Yoruba in Ibadan, Nigeria (West Africa); CHB: Han Chinese in Beijing, China; JPT: Japanese in Tokyo, Japan.

Figure S4

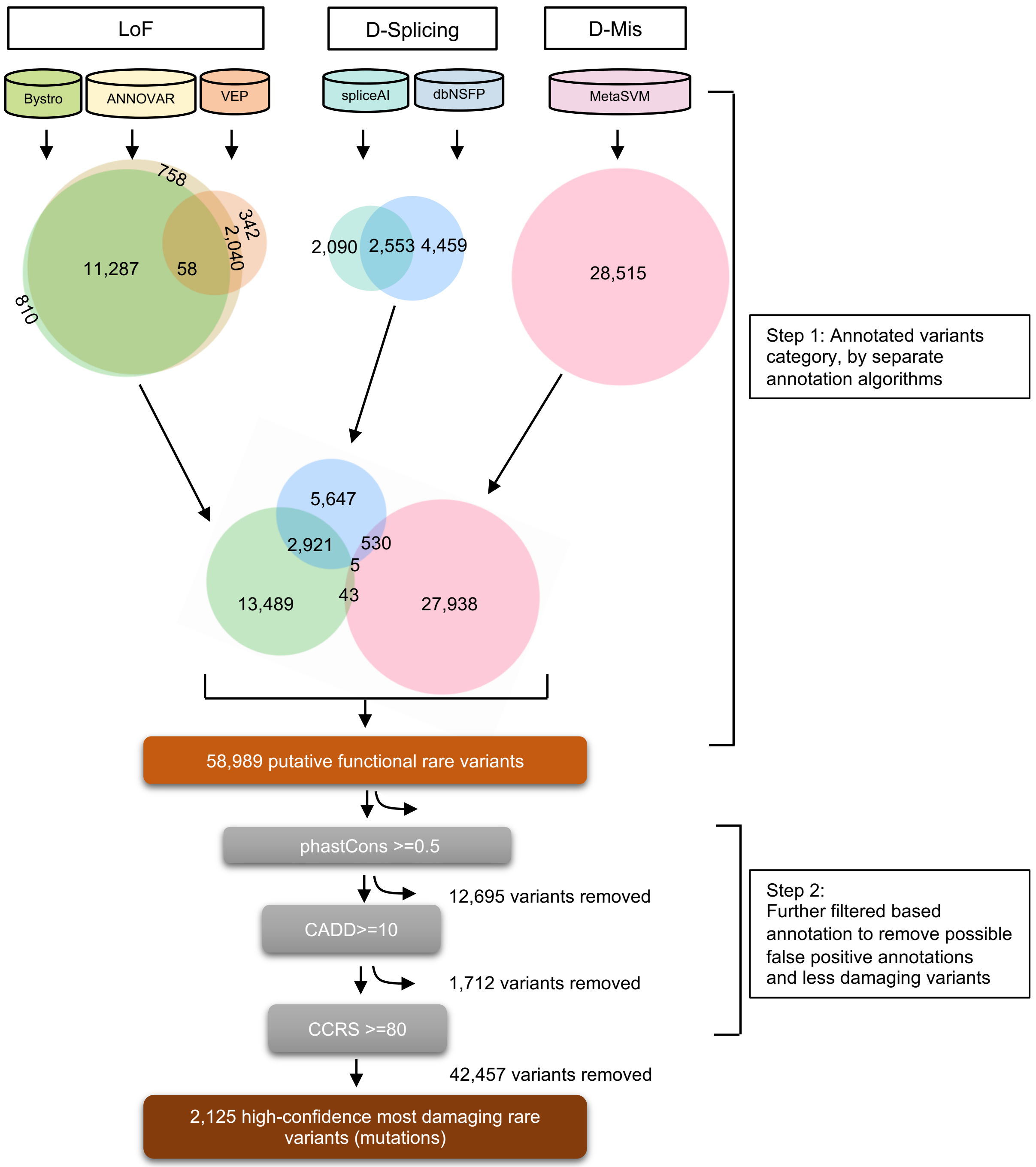


**Figure S4. Identification of 2,125 MDRVs in the cohort from WGS.** Bystro, Annovar and VEP were used to annotate loss of function (LoF) variants in parallel. MetaSVM was used to annotate damaging missense variants (D-Mis). Damaging splicing variants (D-splicing) were annotated by dbNSFP as well as spliceAI. All annotated variants were combined, totaling 58,989 rare putative functional variants and were further filter by phastCons score >=0.5, CADD >=0.5 and CCRS >=80 sequentially to remove false positives. Finally, 2,125 high confidence MDRVs were identified. LoF variants and D-Splicing variants are not mutually exclusive, and subset of the predicted canonical variants were also LoF. Synonymous variants as annotated by Annovar and Bystro are used as negative controls for analyses. CCRS: Constrained coding regions.

Figure S5

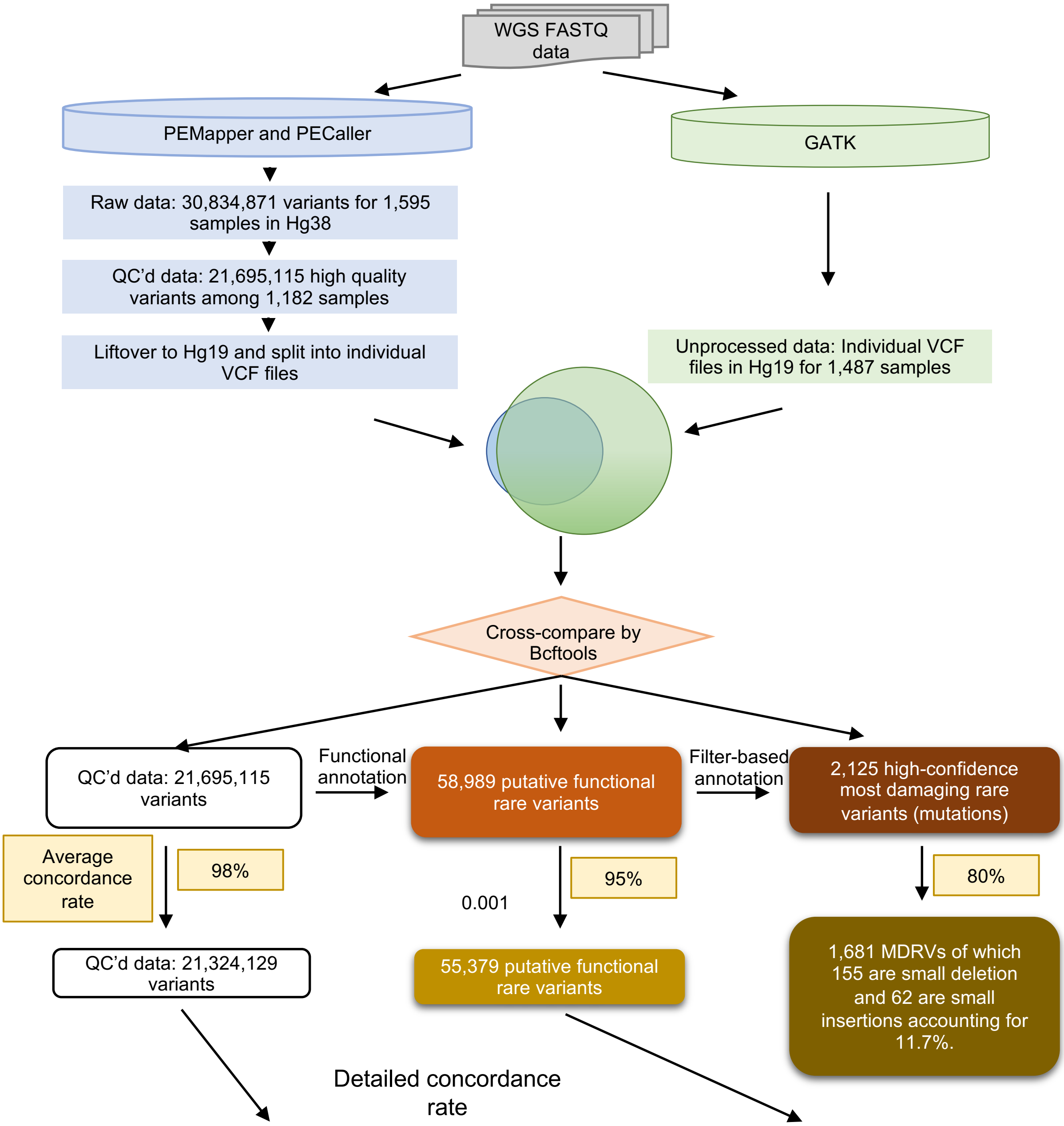


| Concordance rate | All | Indel | SNVs |
|---|---|---|---|
| Consider GT | 0.982±0.004 | 0.833±0.008 | 0.990±0.004 |
| Not consider GT | 0.982±0.001 | 0.841±0.007 | 0.990±0.001 |

| Annotated variants | Number of variants | Concordance rate |
|---|---|---|
| Rare predicted functional variants | SNVs, 48,470 | 96.0% |
| | Indels, 10,519 | 90.6% |
| | Total, 58,989 | 95% |

**Figure S5. Flowchart and results of cross-comparison of variants identified by PEMapper/PECaller within the GATK pipeline.** Of the 21,695,115 quality-controlled high-quality variants, 32,291 failed to liftover to hg19, comprising 0.15% of the total that were cross-compared.

Figure S6

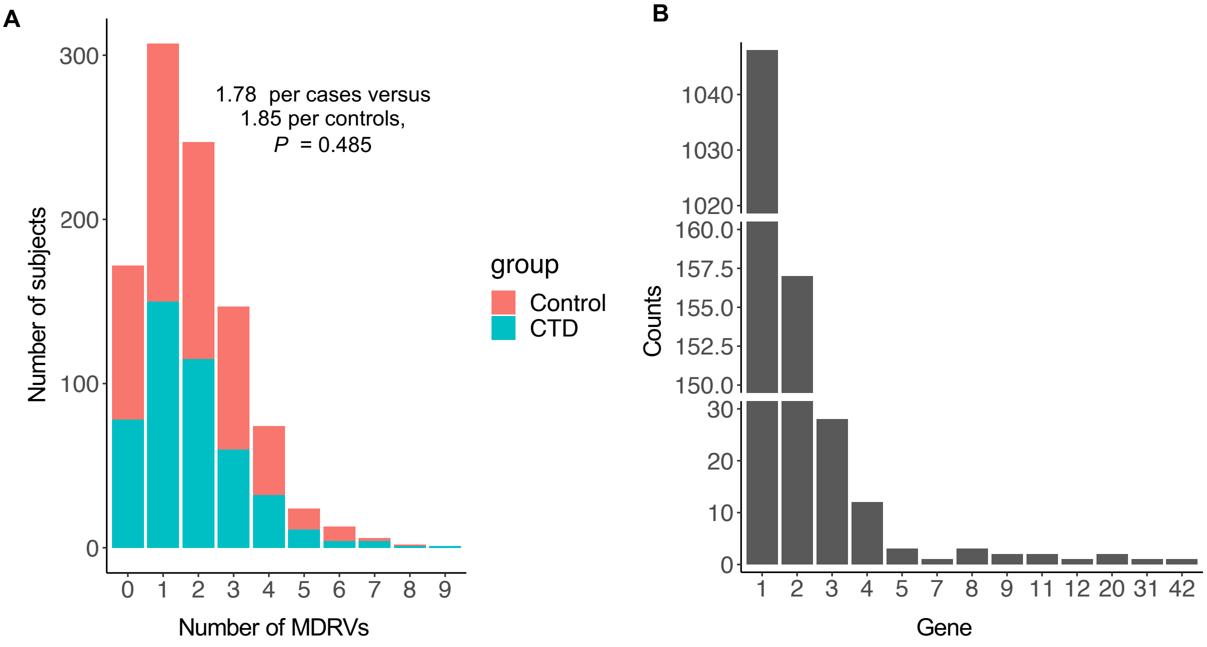


**Figure S6. Distribution of the identified 1,861 MDRVs by case and control status and by gene.** A, Bar plot of distribution of the number of MDRVs (most damaging rare variants) per case and control subject in 22q11.2DS cohort**.** B, Bar plot of the distribution of the number of MDRVs per gene.

Figure S7

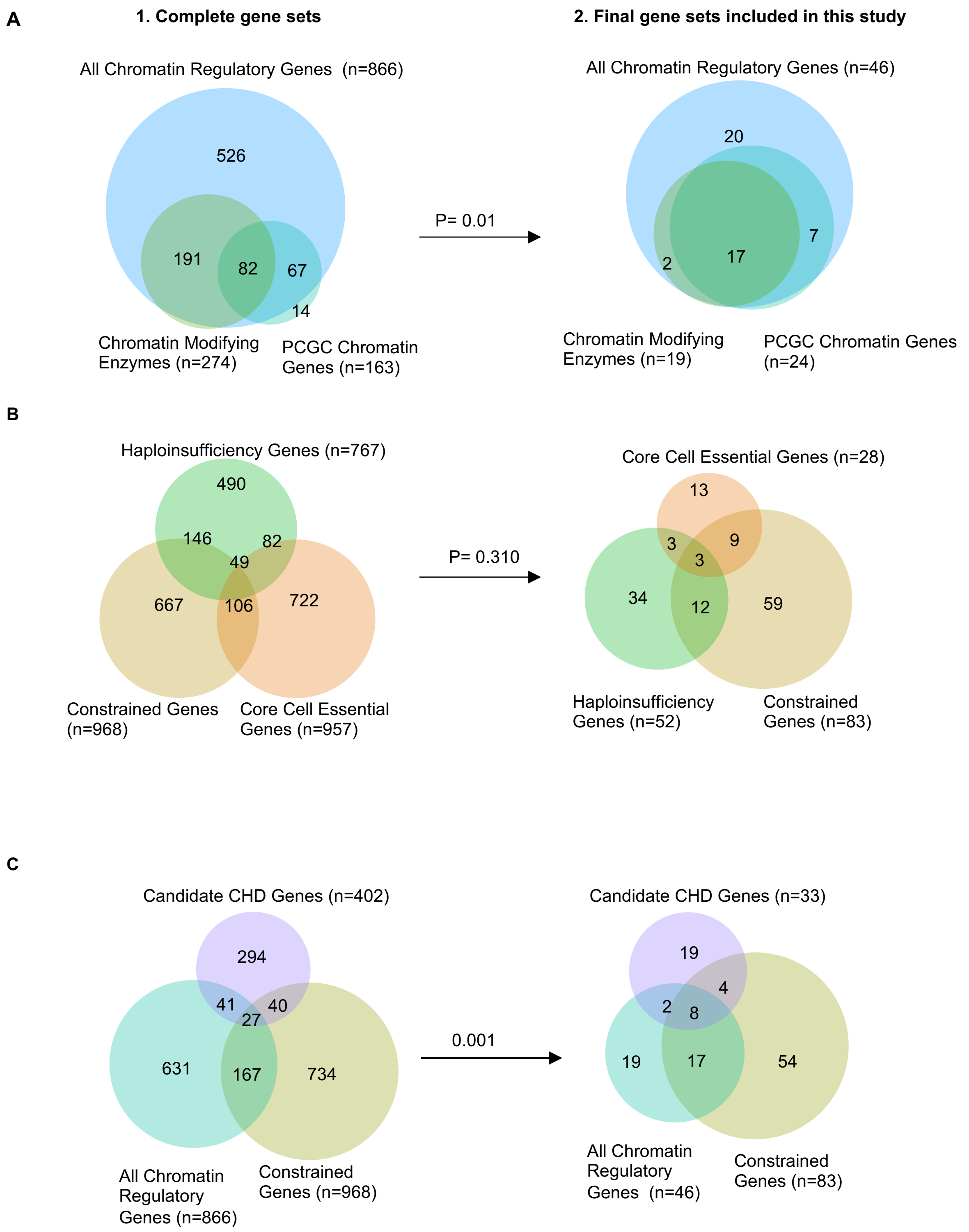
A
1. Complete gene sets
2. Final gene sets included in this study
All Chromatin Regulatory Genes (n=866)
526
191
82
67
14
Chromatin Modifying Enzymes (n=274)
PCGC Chromatin Genes (n=163)
P= 0.01
All Chromatin Regulatory Genes (n=46)
20
7
2
17
Chromatin Modifying Enzymes (n=19)
PCGC Chromatin Genes (n=24)
B
Haploinsufficiency Genes (n=767)
490
146
82
49
667
106
722
Constrained Genes (n=968)
Core Cell Essential Genes (n=957)
P= 0.310
Core Cell Essential Genes (n=28)
13
3
9
3
34
12
59
Haploinsufficiency Genes (n=52)
Constrained Genes (n=83)
C
Candidate CHD Genes (n=402)
294
41
40
27
631
167
734
All Chromatin Regulatory Genes (n=866)
Constrained Genes (n=968)
0.001
Candidate CHD Genes (n=33)
19
4
2
8
19
17
54
All Chromatin Regulatory Genes (n=46)
Constrained Genes (n=83)

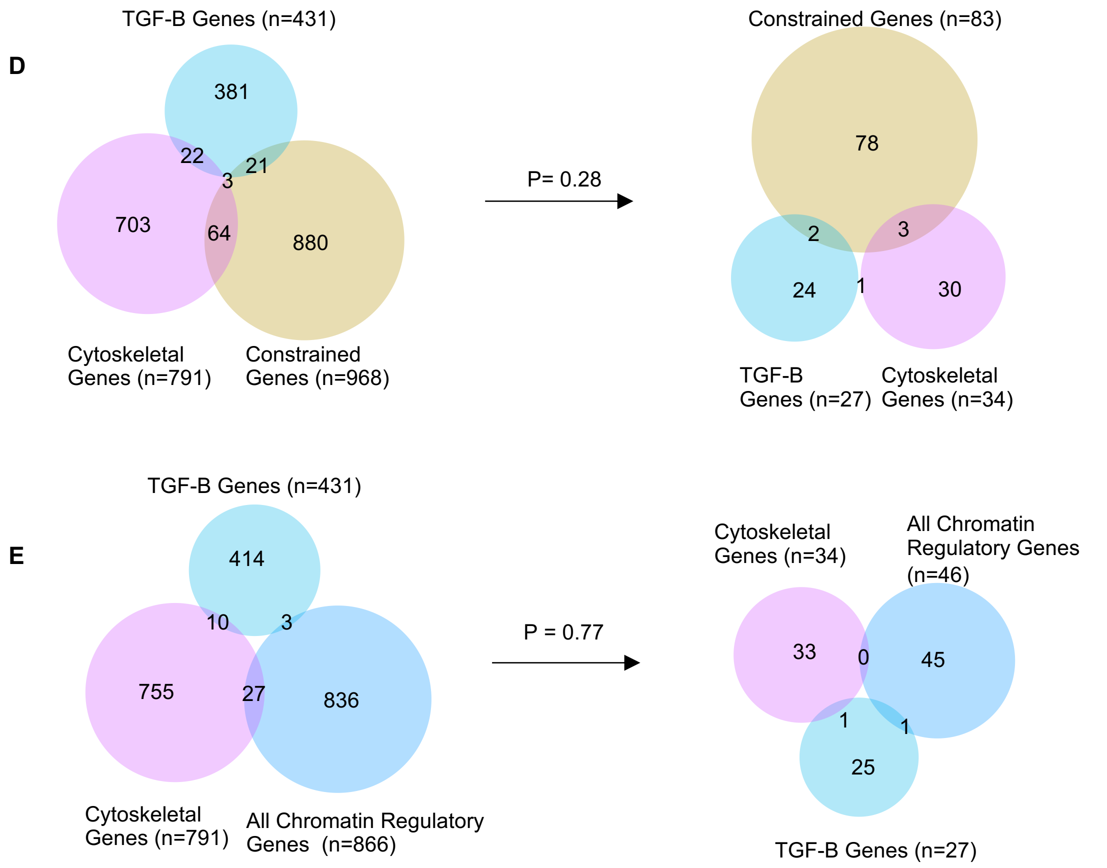


**Figure S7. Overlap of genes within gene sets as well as those of corresponding gene sets for the final number of genes included in the weighted gene set based test.** A, Overlap of the All Chromatin Regulatory Genes set that is compiled from 81 gene set enrichment analysis (GSEA) terms (https://www.gsea-msigdb.org/gsea/index.jsp) (n=866, 46), one gene set was generated using the specific REACTOME term: Chromatin Modifying Enzymes (n=274,19), and 163 Chromatin Genes set was curated by PCGC (n=163, 24). B, Overlap of the Constrained Genes set (n=968, 83), Core Cell Essential Genes set (n=957, 28) and Haploinsufficiency Genes set (n=767,52) for the complete and final gene sets in the cohort. C, Overlap of the genes within the complete and final gene sets of Constrained Genes (n=968, 83), All Chromatin Regulatory Genes (n=866,46) and Candidate CHD Genes (n=402, 33), respectively. D, Overlap of gene sets of Constrained Genes (n=968, 83), TGF-B Genes (n=431, 27) and Cytoskeletal Genes (n=791, 34). E, Overlap of gene sets of Cytoskeletal Genes (n=791,34), All Compiled Chromatin Regulatory Gene (n=866, 46), and TGF-B Genes (n=431,27). P values from the proportion test for the enrichment of the shared genes are indicated.
